# An experimental comparison of the Digital Spatial Profiling and Visium spatial transcriptomics technologies for cancer research

**DOI:** 10.1101/2023.04.06.535805

**Authors:** Taopeng Wang, Kate Harvey, John Reeves, Daniel L. Roden, Nenad Bartonicek, Jessica Yang, Ghamdan Al-Eryani, Dominik Kaczorowski, Chia-Ling Chan, Joseph Powell, Sandra O’Toole, Elgene Lim, Alexander Swarbrick

## Abstract

**Background:** Spatial transcriptomic technologies are powerful tools for resolving the spatial heterogeneity of gene expression in tissue samples. However, little evidence exists on relative strengths and weaknesses of the various available technologies for profiling human tumour tissue. In this study, we aimed to provide an objective assessment of two common spatial transcriptomics platforms, 10X Genomics’ Visium and Nanostring’s GeoMx DSP.

**Method:** The abilities of the DSP and Visium platforms to profile transcriptomic features were compared using matching cell line and primary breast cancer tissue samples. A head-to-head comparison was conducted using data generated from matching samples and synthetic tissue references. Platform specific features were also assessed according to manufacturers’ recommendations to evaluate the optimal usage of the two technologies.

**Results:** We identified substantial variations in assay design between the DSP and Visium assays such as transcriptomic coverage and composition of the transcripts detected. When the data was standardised according to manufacturers’ recommendations, the DSP platform was more sensitive in gene expression detection. However, its specificity was diminished by the presence of non-specific detection. Our results also confirmed the strength and weakness of each platform in characterising spatial transcriptomic features of tissue samples, in particular their application to hypothesis generation versus hypothesis testing.

**Conclusion:** In this study, we share our experience on both DSP and Visium technologies as end users. We hope this can guide future users to choose the most suitable platform for their research. In addition, this dataset can be used as an important resource for the development of new analysis tools.

## Background

Tumours are cellular ecosystems composed of a multitude of cellular subtypes or states. The spatial organisation of cells in tumours is not only a projection of the molecular nature of cancer but also an important predictor for the progression of the tumour and response to treatments [1]. However, previous attempts to characterise the spatial molecular profiles of tumours have been limited by the availability of technology. Conventional spatial molecular technologies, such as multiplexed immunofluorescence, can only examine a handful of markers at a time, restricting our ability to comprehensively map the cellular and molecular features of tumours in tissue [2].

The field of spatial omics technologies has recently expanded rapidly. Novel technologies have encouraged us to revaluate challenges that we were unable to tackle previously. Among these technologies, the GeoMx Digital Spatial Profiling (DSP) platform from Nanostring and the Visium platform from 10X Genomics have emerged as two powerful spatial transcriptomic tools with high data dimensionality and relatively high throughput [2].

DSP is a targeted technology. Instead of profiling the mRNA transcripts themselves, the DSP platform utilises *in situ* hybridisation probes to detect gene expression and later correlate the gene expression profiles with an immunofluorescence image obtained from the same sample [3]. More importantly, the DSP assay allows users to zoom into a specific region of the sample and generate enriched gene expression profiles guided by morphology marker antibodies. This enables deep characterisation of targeted hypotheses in tissues [4].

The Visium assay provides a picture of the global transcriptomic landscape, with a large number (~5000 for Visium slides with 6.5×6.5mm capture areas and ~14000 for slides with 11mm x 11mm capture areas) of densely organised capture spots that are designed to generate a map of gene expression, at relatively high spatial resolution, for evaluation of global localisation and interaction between different cell types [2,5]. It is worth noting that while the Visium frozen tissue assay employs a polyA-based capturing method for fresh frozen samples to directly profile mRNA transcripts, targeted probes are used for formalin-fixed paraffin embedded (FFPE) samples to overcome low RNA quality. Therefore, Visium assays for fresh frozen samples embedded in optimal cutting temperature compound (OCT samples) and FFPE samples should be considered as two independent assays, with potentially different detection capacities.

Previous comparisons between the DSP and Visium platforms generally focused on platform-specific features between the two technologies. These comparisons have comprehensively evaluated the technical specifications provided by the manufacturers and aimed to provide potential users a taste of the best practice for applying these assays to their research [2,6,7,8]. However, given that previous datasets were generated from unmatched samples and often from samples collected under physiological conditions such as normal mouse brain, a thorough comparison on the performance of gene detection and application between the two platform on more challenging sample types such as cancer samples is still missing. In addition, little published data exists for the FFPE Visium assay and how it differs from the OCT Visium assay.

Here, we aim to provide a well-controlled direct comparison between the DSP and Visium technologies. Using preserved cell line samples and primary breast cancer tissue samples, we collected DSP and Visium data from serial sections from the same samples with matched tissue morphologies and cell populations. We started by asking whether there is any fundamental difference between the DSP and Visium platforms in terms of characterising spatial transcriptomic profiles, before going on to assess platform-specific factors. Importantly, we identified several challenges in implementing the DSP and Visium technologies. With these datasets and analyses, we provide a guide to prospective users of these technologies with technical comparisons and insights on experimental workflow design and data processing to assist in decision-making when considering spatial transcriptomic experiments.

## Methods

### Sample collection

Surgical specimens were assessed and sampled by a pathologist with specialist experience in breast cancer to ensure the tumour area was collected. Tumour samples were trimmed of excess fat and macroscopic necrosis and then cut to size. For samples 4747, 4754 and 4806, tissues were sliced in the middle to form two pieces with mirrored morphology and preserved as FFPE or OCT samples respectively. For sample 4766, the tissue was thin and only adjacent pieces were preserved as FFPE or OCT samples. For FFPE samples, tissues were fixed in 10% neutral buffered formalin (NBF) for 24 hours before changing to 70% ethanol prior to processing and embedding. For OCT samples, tissues were placed mirrored face-down onto a flat metal spatula. Tissues were submerged in an isopentane bath in a metal beaker surrounded by dry ice until frozen through. Frozen tissues were gently removed from the spatula and submerged into precooled OCT and then frozen by surrounding the mould with crushed dry ice. All OCT tissue samples have a RIN value of at least 7.

To create ‘synthetic tissue’ references, cultured Jurkat and SKBR3 cells were collected as single-cell suspension and washed twice using 1x PBS. Jurkat and SKBR3 cells were then counted and mixed at 6 different ratios (0:100, 5:95, 30:70, 70:30, 95:5 and 100:0) to create a gradient. The prepared cell mixes were then converted into OCT or FFPE blocks. For OCT cell array samples, an OCT mould was made by incubating OCT with 6 small metal pillars on dry ice. The OCT were given time to solidify but not fully set to avoid attachment to the metal pillars. Once the metal pillars were removed, the OCT mould was given extra time to fully solidify resulting in 6 holes in the mould. The mixed Jurkat and SKBR3 cells were then pelleted at 300g for 5min at 4 degrees and the supernatant was removed. The rest of the cells and buffer were then mixed by gentle flicking. 10µl of the cells from each mixing ratio was transferred to a corresponding hole in the OCT mould and frozen on dry ice. For FFPE cell array samples, mixed Jurkat and SKBR3 cells were firstly resuspended in 10% NBF. The resuspended cells were then pelleted immediately at 700g for 10min to remove the NBF in the supernatant. The cell pellets were then mixed with equal volume of warm 3% agarose and transferred to the lids of PCR tubes. Once solidified, the cell pellet samples were retrieved from the lids using a metal scalpel with care. The resulting cell pellets were stored in 70% ethanol for 24 hours before being processed into FFPE blocks.

Before sectioning for gene expression experiments, an H&E stained section was obtained from each FFPE or OCT sample to evaluate the morphological features of the sample and to guide the selection of DSP regions of interest (ROIs). For both DSP and Visium assays involving FFPE samples, closest possible 5μm thick sections were used. For DSP experiments, the first 2 sections were discarded before collecting the samples for DSP experiments. The prepared sections were stored at −20°C. For OCT samples, serial sections were prepared at the thickness of 7 or 10μm for the DSP or Visium experiments respectively. 10μm thick sections were used for Visium tissue optimisation experiments.

### Nanostring morphology marker antibody conjugation

CD8 antibody (Clone AMC908, ThermoFisher) was conjugated using Alexa Fluor™ 647 Antibody Labelling Kit (ThermoFisher). 100mg of CD8 antibody at a concentration of 1mg/mL was cleaned twice using the Zeba™ Spin Desalting Columns, 7K MWCO (ThermoFisher). The filtered antibody was then mixed with 1M sodium bicarbonate (pH 8.5) at a ratio of 10:1 by volume. The primed antibody was then transferred immediately to the tube with fluorescent dye supplied by the kit. The antibody-fluorophore mixture was homogenised by pipetting and incubated at room temperature for 1 hour in dark. During incubation, the antibody-fluorophore mix was mixed every 15min by gentle flaking on the tube. After incubation, the antibody-fluorophore mix was filtered twice with the Zeba Spin columns. The conjugated antibody was stored at 4°C until use.

### DSP experiment

DSP experiments were conducted according to manufacturer’s instructions (Slide prep manual version: MAN-10115-04; DSP instrument operation manual version: MAN-10116-04; Library prep manual version: MAN-10117-04) with minor adjustments. Briefly, sectioned FFPE slides were firstly baked for 60 minutes at 65°C followed by dewaxing and antigen retrieval. For OCT sections, samples were firstly thawed and fixed in 10% NBF for 16 hours at room temperature. The fixed samples were then washed 3 times in 1x PBS followed by antigen retrieval. From antigen retrieval, the FFPE and OCT samples were treated with the same conditions. Both FFPE and OCT cell array samples were incubated for 5min at 100°C in 1x Antigen Retrieval Solution (ThermoFisher, 00-4956-58) using a pressure cooker. For tissue samples, samples were incubated for 20min at the same temperature. For proteinase K digestion, samples were incubated with 0.1μg/mL proteinase K (ThermoFisher, AM2546) in a in a 37°C water bath. The cell array samples were incubated for 5min while the tissue samples were digested for 15min. For in situ hybridisation, 240μL of diluted DSP probe mix was added to each slide and incubated at 37°C in a hybridisation oven for 18 hours over night. The processed slides were then labelled with morphology marker antibodies. For cell array samples, SYTO13, anti-pan-cytokeratin-AF532 antibody (Nanostring) and anti-CD45-AF594 antibody (BioLegend, 103144) were used. For tissue samples, SYTO13, anti-pan-cytokeratin antibody and the anti-CD45 antibody from the Nanostring solid tumour morphology marker kit were used in conjugation with the conjugated anti-CD8 antibody as mentioned above to illustrate tissue morphology. Areas of illumination (AOIs) were collected using a range of shapes and sizes, as appropriate to the experiment. In some cases, to allow direct comparison with the Visium platform, ~55μm diameter non-segmented AOIs were captured to ‘mimic’ the data generated by the Visium platform.

After DSP collection, samples were dehydrated at 65°C for 1.5 hours in a thermo-cycler with the lid kept open. Samples were then rehydrated and subjected to sequencing library preparation. During the experiment, we noticed that the volume of the primers in wells A1, H1, A12 and H12 from SeqCode plate B, well E1 from SeqCode plate F and well H1 from SeqCode plate G was lower than the volume in the other wells. Instead, libraries for samples in these wells were synthesised using primer from well C1-C6 from SeqCode plate H. After library synthesis, 4μL of PCR products from each well were firstly pooled together by type i.e. Visium-mimic AOIs, segmented AOIs, size gradation AOIs and biological AOIs (Fig. 1c-d). The resulting pools of PCR products were then merged together adjusting for the total area size of all AOIs in each pool to guarantee a comprehensive sampling of smaller AOIs. The pooled sequencing library was then quality controlled and sequenced on a NovaSeq 6000 instrument (Illumina). Paired-end and dual-indexed reads were generated in the format of 2 x 28bp with an additional 2 x 8bp for index sequences.

**Figure 1:**
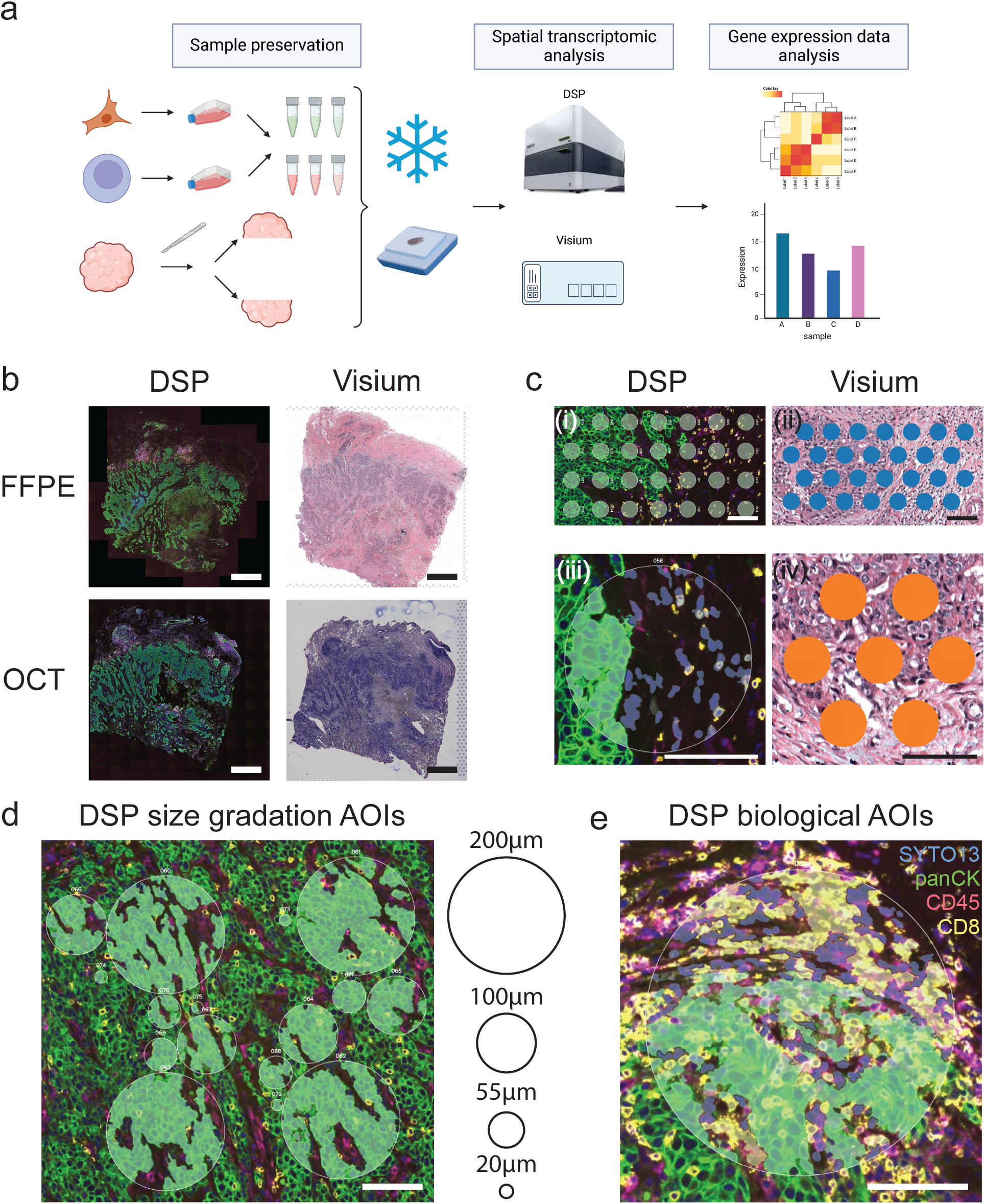
Experiment overview. (**a**) Schematic illustration of sample preservation and experiment workflow. Cultured Jurkat and SKBR3 cells were mixed at six different ratios and preserved as OCT or FFPE samples. Tissue samples were sliced in the middle and the resulting two pieces were preserved as OCT or FFPE samples respectively. Closest possible sections were used for DSP and Visium assays. The illustration was created with BioRender.com. (**b**) Example images of tissue morphology. FFPE and OCT sections from sample 4806 were processed for DSP or Visium assays. Note the overall matching morphology between FFPE and OCT samples and between sections used for DSP and Visium assays. Scale bars = 1mm. (c) Example of DSP AOIs and Visium spots used for direct comparison. (**i**) Example of Visium mimic AOIs across the tumour-stroma interface. (**ii**) Example of Visium spots across the tumour-stroma interface used for direct DSP and Visium comparison. (**iii**) Example of segmented comparison DSP AOIs. Each AOI was segmented into pan-cytokeratin positive and pan-cytokeratin negative segments according to the immunofluorescence signal. **(iv**) Example of Visium spots located in the matching region where segmented comparison DSP AOIs were collected. Scale bars = 100μm. (**d**) Example of size gradation and biological DSP AOIs to test the performance of segmentation of the DSP platform. Scale bars = 100μm.

RNAse free or buffer was used throughout the experiment except for xylene and ethanol for histology. All surfaces were decontaminated using RNAse ZAP.

### Visium experiment

OCT Visium experiments were conducted according to the manufacturer’s protocol. The optimal tissue permeabilisation time for each sample was determined using the Visium tissue optimisation kit. The resulting RNA footprint fluorescence images were reviewed and 17, 16 and 12 min were used for gene expression experiments for the cell array sample, 4747, 4754 and 4806 respectively. The rest of the procedures were conducted according to the Visium protocol. The generated cDNA library was then sequenced on a NovaSeq 6000 instrument (Illumina).

FFPE Visium data was generated by 10X Genomics with no deviation from the protocol. The H&E images were taken using a Metafer slide scanning system (Metasystems) with a Zeiss Plan-Apochromat 10x/NA 0.45 objective lens. The resulting cDNA libraries were sequenced on a NovaSeq 6000 system (Illumina).

### FASTQ file processing

For Visium data, demultiplexed FASTQ files were converted to count matrices using SpaceRanger 1.3.1 (10X Genomics). OCT and FFPE Visium data were mapped to refdata-gex-GRCh38-2020-A (10X Genomics). Visium Human Transcriptome Probe Set v1.0 GRCh38-2020-A (10X Genomics) was also provided for processing the FFPE Visium data. Spot annotation was conducted in loupe browser 5 (10X Genomics). A Seurat object (Seurat V4) was then constructed for each sample using deduplicated count matrices and spot annotations [9].

For DSP data, raw reads from FASTQ files were mapped to the Hs_R_NGS_WTA_v1.0 reference (Nanostring) using the GeoMx NGS pipeline software V2.2 (Nanostring) and saved as Digital Count Conversion (DCC) files. The DCC files were then used to construct a gene expression count matrix using the GeomxTools package V3.0.1 [10].

DSP and Visium data was also down sampled to account for technical variations in direct DSP and Visium comparison. FASTQ files from samples used for direct DSP and Visium comparison were selected and down sampled in a per sample manner. Raw reads were down sampled so as to not exceed the minimum recommended levels (100 reads/µm^2^ for DSP; 25,000 reads/spot for FFPE Visium; and 50,000 reads/spot for OCT Visium) for each platform using seqtk 1.3 [11]. For samples whose sequencing depths are below the recommendations, raw data was kept as is. The resulting down sampled FASTQ files were aligned to the references using the same tools and references as mentioned above.

### Data quality control (QC) and filtering

For Visium data, spots with less than 1000 unique molecular identifiers (UMIs) detected were considered low-quality and excluded from the data. In addition, spots underneath regions with tissue processing artefacts were manually annotated and excluded as well.

For DSP data, data were quality controlled per individual AOI. AOIs were excluded from the dataset if they met any of the following conditions: less than 80% of reads aligned to the reference, less than 40% sequencing saturation, or less than 1000 UMI. After QC and filtering, DSP count matrix and annotation were saved as Seurat objects for more consistent accessing and analysis.

### Data normalisation and differential expression (DE) analysis

For gene expression visualisation, DSP and Visium data were normalised respectively using the “NormalizeData” function with default settings in the Seurat package. For DE analysis in DSP and Visium comparison, DSP visium mimic AOIs and Visium spots from matching regions were selected. For Visium data, data was normalised on a per sample basis using the same “NormalizeData” function in the Seurat package. For DSP data, AOIs collected from the cell array samples and tissue samples were grouped separately to minimise the impact of tissue composition on data normalisation. AOIs in each group were then normalised using the Q3 method per manufacturer’s recommendation. Briefly, a 3^rd^ quantile threshold was calculated for each AOI for the estimation of normalisation factors across AOIs within the same group. The data was then normalised using the normalisation factors. Both DSP and Visium data was filtered for outlier genes before DE analysis. For Visium data, genes detected with equal or more than 1 count in at least 3 technical replicates were kept for DE analysis. For DSP data, a limit of quantification (LOQ) was estimated for each AOI. The LOQ was calculated using the following formula with raw counts of negative control probes:

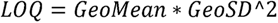

Only genes with expression above LOQ in at least 3 technical replicates were included in the DE analysis. The normalised and filtered DSP and Visium data was then fitted to a linear model on a per sample basis using the limma package (version 3.52.2) and t-statistics were calculated using the “eBayes” function in limma [12].

For DE analysis between CD8 AOIs and non-CD8 AOIs, DSP data was grouped according to pathology annotation. DE analysis was conducted in a similar way as mentioned above.

Genes with an adjusted P value (Benjamini-Hochberg method) less than 0.05 were considered to be differentially expressed. DE results were visualised using barplots (ggplot), correlation heatmap (ComplexHeatmap v2.12.0) [13]. Top DEGs were selected based on the absolute value of the t statistics in DE analysis results.

### Cell type deconvolution

The cellular composition of each Visium spot and DSP AOI was predicted using published single-cell transcriptomic signatures [5]. Both the “major” and “minor” level cell type signatures were used for deconvolution.

For Visium data, cell type deconvolution was conducted using Stereoscope v0.3.1 [14] based on the top 2000 highly variable genes as defined by scanpy v1.7.2 [15] using the “Seurat” flavoured method [16].

For DSP data, deconvolution was conducted using all genes with the SpatialDecon package [17]. The Q3 normalised, log2-scaled gene expression matrix was used as the input for the analysis. All other parameters were kept as default.

### Pre-ranked GSEA analysis

Genes were ranked based on t statistics from DE analysis. Pre-ranked GSEA analysis was then conducted using the GSEA software (linux, v4.2.2) [18,19]. Genes were mapped to gene ontology biological processes pathways (v7.5.1) obtained from the Molecular Signatures Database (MSigDB)[20]. The results of pathway analysis were then visualised using the ComplexHeatmap package in R.

### Visium data dimension reduction, clustering and over-representation analysis (ORA)

Dimensionality reduction was conducted on Visium data for each individual sample using the Seurat package. Original datasets (without down-sampling) were used for this analysis. Gene expression data was normalised using the ScTransform method (V1) [21]. Dimension reduction was conducted using PCA and UMAP methods. A total of 30 principal components were used for dimension reduction through UMAP method. Visium spots were then clustered using the Seurat package. The optimal clustering resolution was selected based on the spatial distribution of common cell type marker genes. The clustree package (v0.5.0) was also used to evaluate the relationship between clusters at different clustering resolution [22]. The clustering resolution that provides relatively high clustering stability but also reflects the biological complexity of the tissue was selected as the optimal clustering resolution for each sample.

Clusters predominantly containing cancer cells were identified based on the proportion of cancer cells predicated by deconvolution as well as the tissue morphology underneath the spots in each cluster. Differential gene expression analysis between the identified cancer clusters was then conducted using the “FindAllMarkers” function in the Seurat package. Default parameters for Seurat v4 were used. All genes with an adjusted P value less than 0.05 were considered significantly differentially expressed and passed to the downstream ORA analysis using the clusterProfiler package (v4.4.4) [23]. Enriched hallmark pathways were calculated using “compareCluster”. The top 10 pathways upregulated in each cluster were then visualised using the “dotplot” function in the clusterProfiler package.

## Results

### Experimental design

In total, 4 primary breast cancer tissue samples and 2 cultured cell lines were preserved for DSP and Visium comparison. Patient 4747 was diagnosed with ER+ breast cancer while 4754, 4766 and 4806 were all diagnosed with triple negative breast cancer (TNBC) by clinical examination. All tissue samples were sliced in the middle to form 2 pieces of tissue with mirrored morphology except for 4766 from which adjacent pieces were taken (Fig. 1a). The resulting two pieces were then preserved as FFPE or OCT blocks respectively (Fig. 1b). To permit more direct quantitative comparisons between platforms, cultured cell lines were mixed to create ‘synthetic tissues’ with controlled cellular proportions. We selected Jurkat (T lymphoma) and SKBR3 (breast cancer) cell lines as representations of immune and epithelial malignant cell types, respectively. Jurkat and SKBR3 cells were mixed at 6 ratios: 100-0, 95-5, 70-30, 30-70, 5-95 and 0-100. The mixed cell samples were then divided into two aliquots for FFPE and OCT sample preservation. Cell samples with different mixing ratios were arrayed together to generate a cell microarray block for FFPE or OCT samples respectively. Serial sections were then cut for DSP and Visium assays.

The DSP and Visium platforms collect spatial transcriptomic data in a very different manner. The Visium platform generates a uniformed array of ~5000-14000 spots per sample depending on the size of the capture area, while the DSP AOIs are manually selected in locations of interest and can have different sizes or segmentation based on morphology and marker expression. To facilitate a direct comparison between DSP and Visium assays, 4 types of DSP AOIs were collected: 1) Visium-mimic AOIs, 2) segmented AOIs, 3) size gradation AOIs, and 4) biological AOIs. These AOI types are defined as follows. 1) Visium-mimic AOIs are circular AOIs 55μm in diameter aiming to mimic data collection of visium spots. 4 Visium mimic AOIs were selected in each cell pellet with a different SKBR3 / Jurkat mixing ratio, and 28 Visium-mimic AOIs were selected in tissue samples 4747, 4754 and 4806 (Fig. 1c i-ii). 2) Segmented AOIs are circular AOIs, 200μm in diameter, segmented into epithelial and non-epithelial compartments using immunofluorescence-guided masks (Fig. 1c iii). 3) Size gradation AOIs are circular AOIs varying in size from 20μm in diameter to 200μm in diameter and used to evaluate the impact of AOI size on DSP transcriptomic data collected (Fig. 1d). 4) “Biological AOIs” are circular AOIs collected around biological structures annotated by pathology, such as immune clusters adjacent to the tumour. Depending on the cellular composition in each location, epithelial, non-epithelial and CD8 segments were collected (Fig. 1e).

### Comparison of the transcriptomic coverage and detection sensitivity of the DSP and Visium platforms

To compare the performance of the DSP and the Visium platforms under more compatible conditions, we evaluated the impact of two major technical factors on the performance of the two platforms: AOI size (DSP) and sequencing depth.

For DSP assays, the size of each AOI can be manually adjusted, allowing sampling of different numbers of cells per AOI. To test the impact of AOI size, we focused on size gradation AOIs (Fig. 1d). In line with the previous literature [3,24], we observed a positive correlation between the size of AOI and the number of genes with at least 1 UMI detected per AOI (Fig. S1). While more than 5000 genes were detected in 20um spots, sensitivity increased markedly between the 20um and 55um spot size. Therefore we mainly focused on 55um AOIs that are of comparable size to Visium spots (Fig. 1c i-ii).

Sequencing comprises a substantial component of the total cost in spatial transcriptomics, and sequencing depth affects sensitivity of detection [25]. To address the optimal sequencing needs of each platform, we firstly compared the performance of each platform as a function of sequencing depth. For DSP WTA assays, a minimum of 100 reads per μm^2^ is recommended [26]. Most of the DSP samples processed were able to reach and surpass this threshold (Fig. S2a). We did observe a few outlier samples. However, the sequencing saturation of all DSP samples have surpassed 50% indicating proper profiling of the sequencing libraries (Fig. S2c). On the other hand, a minimum of 25,000 or 50,000 reads per spot was recommended for the FFPE and OCT Visium assays, respectively [27,28]. The current FFPE Visium datasets were extensively sequenced exceeding the threshold by at least 1-fold (Fig. S2b). Samples processed using the OCT Visium assays were at or slightly below the required sequencing depth (Fig. S2b). Interestingly, while the FFPE Visium samples were sequenced deeper as compared to OCT Visium samples, we observed an inverse trend in sequencing saturation (Fig. S2c), indicating a higher library diversity of the FFPE Visium samples as compared to the OCT Visium samples.

To account for the variations in sequencing depth between DSP and Visium, as well as between individual samples processed using the same platform, we down-sampled the gene expression data from AOIs/spots used for direct comparison to the recommended read depths at a per-sample level (Fig. S2d-f). Datasets already below the recommendations were kept as is (Fig. S2d-f). As expected, we observed a decrease in sequencing saturation for all samples after down-sampling. Most impacted was the FFPE Visium data which exceeded the recommendation by the greatest extent. All FFPE Visium samples only achieved around 10-20% sequencing saturation after down-sampling, whereas minimal impact was observed for DSP and OCT Visium data (Fig. S2f). This suggests that sequencing Visium FFPE libraries above the recommended depth is necessary to achieve saturation >50% in human cancer studies.

Using these standardised gene expression datasets, containing the same number of DSP Visium-mimic AOIs and Visium spots, we examined transcriptomic coverage provided by the DSP and Visium assays. The biotype of transcripts detected were annotated using the GRCh38 reference and briefly summarised into 5 groups: 1) mitochondrial RNA (MT); 2) RNA for ribosomal proteins (RP); 3) RNA for T cell receptors (TCR) or B cell receptors (BCR); 4) RNA for other proteins and 5) non-coding RNA (ncRNA). As expected, the OCT Visium assay is the only assay detecting mitochondrial RNA and the main assay detecting non-coding RNA due to the non-targeted capturing using poly(T) capture handles (Fig. 2a). The DSP platform contains probes targeting genes coding for ribosomal proteins (see column “DSP_panel”, Fig. 2a), which are also detected by the OCT Visium, but mostly absent from the FFPE Visium probe-set. In contrast, the FFPE Visium assay includes many more probes against TCRs and BCR gene segments than the standard DSP probe set (Fig. 2a), which will be valuable in the investigation of tumour immunology.

**Figure 2:**
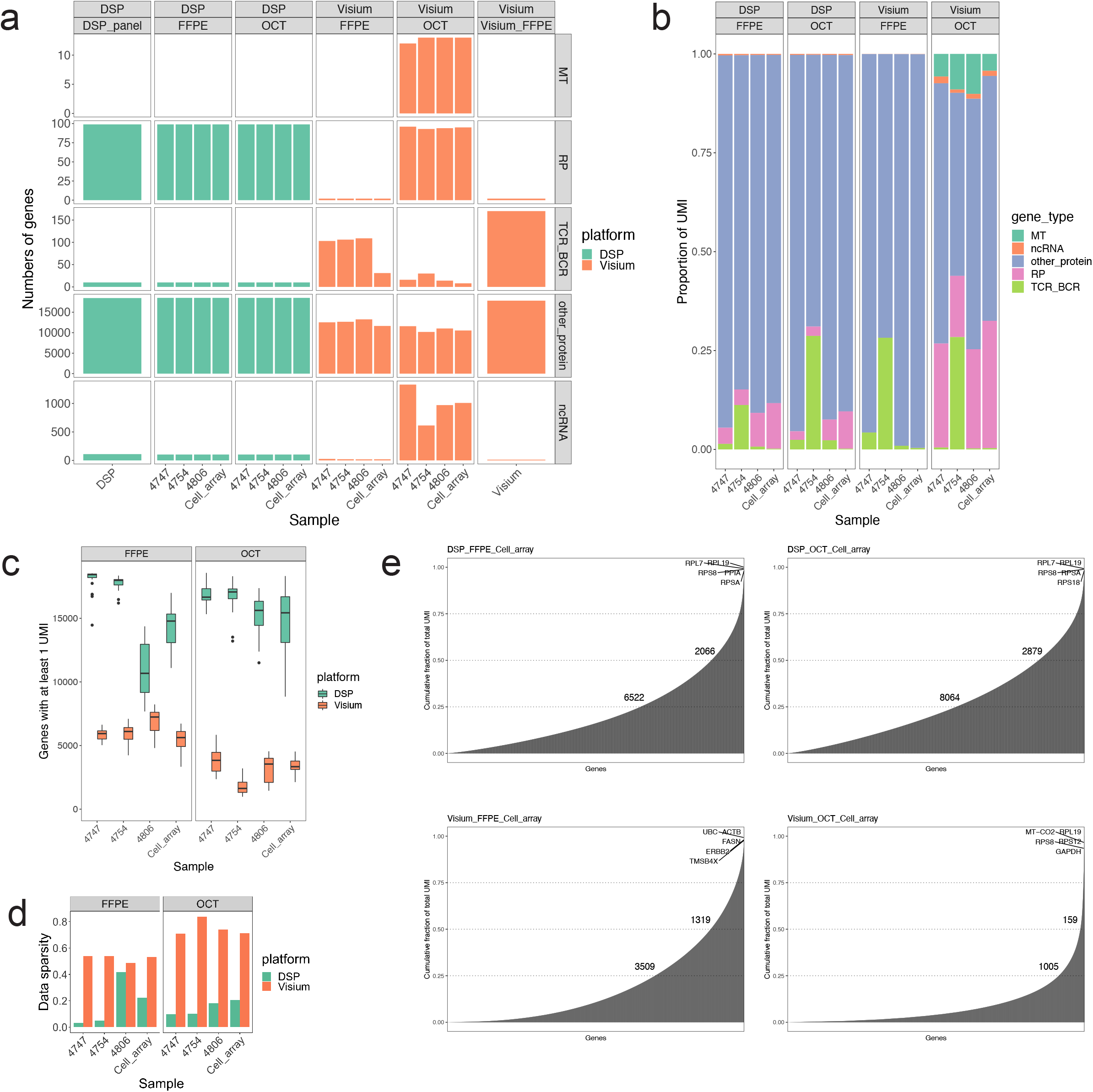
Comparison of the DSP and Visium platform on transcriptomic coverage and sensitivity. Visium mimic AOIs in DSP data and Visium spots collected from the same regions were used for the analysis (**a**) Barplots illustrating the types of genes with at least 1 UMI detected by the DSP and Visium assays. Probes included in the whole DSP and FFPE Visium panels were also plotted for comparison (“DSP_panel” & “Visium_FFPE”). (**b**) Barplots illustrating the proportion of counts for each type of genes in each sample detected by the DSP and Visium assays. (**c**) Boxplots illustrating the numbers of genes with at least 1 UMI detected per AOI or spot in DSP or Visium data. (**d**) Barplots illustrating the sparsity of the gene expression matrix. The sparsity was calculated as the proportion of zero counts in the matrix. (**e**) Barplots illustrating the ranking of genes and their contribution to the total counts collected in each dataset.

We also evaluated the number of molecules, also known as unique molecular indices (UMIs), from each type of RNA transcript detected by the DSP and Visium platforms. The majority of the counts in each assay were related to protein-coding genes (Fig. 2b). Around 30-40% of UMIs collected by the OCT Visium assay were related to transcripts for mitochondrial or ribosomal proteins. In addition, all assays seem to detect substantial amount of UMIs in samples from patient 4754 for TCR or BCR transcripts, potentially reflecting variations in tissue immune cell composition between the samples (Fig. 2b).

The results above suggested that both the DSP and Visium platform can provide an overall good transcriptomic coverage of the samples profiled, but that coverage for specific applications varies by platform. We next aimed to evaluate the sensitivity of gene expression detection per spot level. Using genes with at least 1 count detected as threshold, the DSP assays in general detected many more genes per spot than the Visium assays (Fig. 2c). As a consequence, the DSP data had less zero observations in the gene expression matrix as reflected by the overall low matrix sparsity (Fig. 2d). In line with this observation, UMIs are more evenly distributed across genes in DSP assays with ~6000 – 10000 genes contributing to 75% of all UMIs collected (Fig. 2e; Fig. S3). On the other hand, the counts collected in Visium datasets are concentrated among a smaller group of genes with ~3000 and ~1000 genes occupying 75% of all UMIs in the FFPE or OCT Visium data, respectively (Fig. 2e; Fig. S3). Therefore, these results suggest that the DSP assays are more sensitive than the Visium assays given more genes were detected with counts and the UMI distribution is more even across the transcriptome.

However, DSP assays are known to contain noise due to non-specific probe binding [3]. Non-targeting control probes are included in the probe panel in order to model the level of non-specific binding in each AOI. A LOQ threshold is often applied to evaluate if a gene is considered to be detected or not in the DSP datasets. Genes recurrently below the LOQ threshold can then be excluded from analysis to highlight the key biology of the samples [3]. Targeted probes were also used in the FFPE Visium assays. However, the probes are designed to contain a left-hand side and a right-hand side so that only reads from both probe pairs are included in the final count matrix. This potentially allows the exclusion of some non-specific readings from the dataset. In contrast, the Visium OCT assay employs an unbiased polyA-based capturing approach and is free from the bias due to variations in probe sequences.

We then evaluated the specificity of detection in both DSP and Visium data. Given that the cell array samples only contain Jurkat (T-cell lymphoma) and SKBR3 (breast cancer) cell-lines, we reasoned that immunoglobulin heavy chain genes should not be detected in this sample. Indeed, no reads from immunoglobulin heavy chain genes were detected by the FFPE or OCT Visium assays (Fig. S4a). However, unfiltered DSP data did contain non-specific readings for immunoglobulin heavy chain genes. These results are in line with previous studies, which showed the presence of non-specific signals in the DSP assays [3]. Non-specific detection in the DSP data can be reduced by filtering the count matrix with the geometric mean of non-target probe readings (Fig. S4b) and was completely removed by using LOQ filtering (Fig. S4c). However, LOQ filtering may also introduce false negatives. For example, EPCAM is a well-established epithelial cell marker. Filtering of DSP gene expression data using the LOQ method can lead to exclusion of EPCAM signal in several AOIs containing almost exclusively SKBR3 cells (Fig. S4c). In addition, we noticed that the sensitivity of the DSP platform drops after applying additional filtering. For example, when the raw counts were filtered using the geometric mean of non-target probe readings, we noticed a clear decrease in the numbers of genes detected per spot (Fig. S5a). Similarly, the sparsity of the gene expression matrix also increased (Fig. S5b; Fig. 2d). Interestingly, the numbers of genes contributing to majority of the UMIs collected (75%) in background filtered DSP data are comparable to that of in the FFPE Visium assay (Fig. S5c-j). Non-surprisingly, the sensitivity of the DSP assays can drop even further after more stringent filtering is applied (LOQ filtering) (Fig. S6).

### Comparison of the DSP and Visium platforms on detecting gene expression changes

A major application of spatial transcriptomics platforms is in the determination of the difference in gene expression between cellular compartments, so we asked how well the DSP and Visium platform capture biological variation within samples. Using the cell-pellet datasets, we first evaluated the expression of marker genes in DSP and Visium data. We observed high expression of epithelial markers such as KRT18 in cell samples containing higher proportions of SKBR3 cells and vice versa for immune markers in samples with more Jurkat cells, thereby showing good correlation between gene expression and cellular composition in all datasets (Fig. 3a; Fig. S7). Turning to tissue samples, a similar trend was observed in most of the datasets compared. For both FFPE and OCT DSP data, there are clear differences in epithelial or TME marker gene expression between regions with high epithelial content and regions with low epithelial content (Fig. 3b; Fig. S8). For FFPE Visium data, the trend is generally clear for sample 4754 and 4806 but less so for 4747. While in line with the DSP data showing a significant enrichment of KRT18 expression in the ‘high epithelial’ region as compared to the ‘low epithelial’ region in 4747 (Fig. 3b), no enrichment was observed for KRT8 in FFPE Visium data (Fig. S8). Also, the difference in marker gene expression between regions with high or low epithelial content detected by the OCT Visium assay is small (Fig. 3b; Fig. S8).

**Figure 3:**
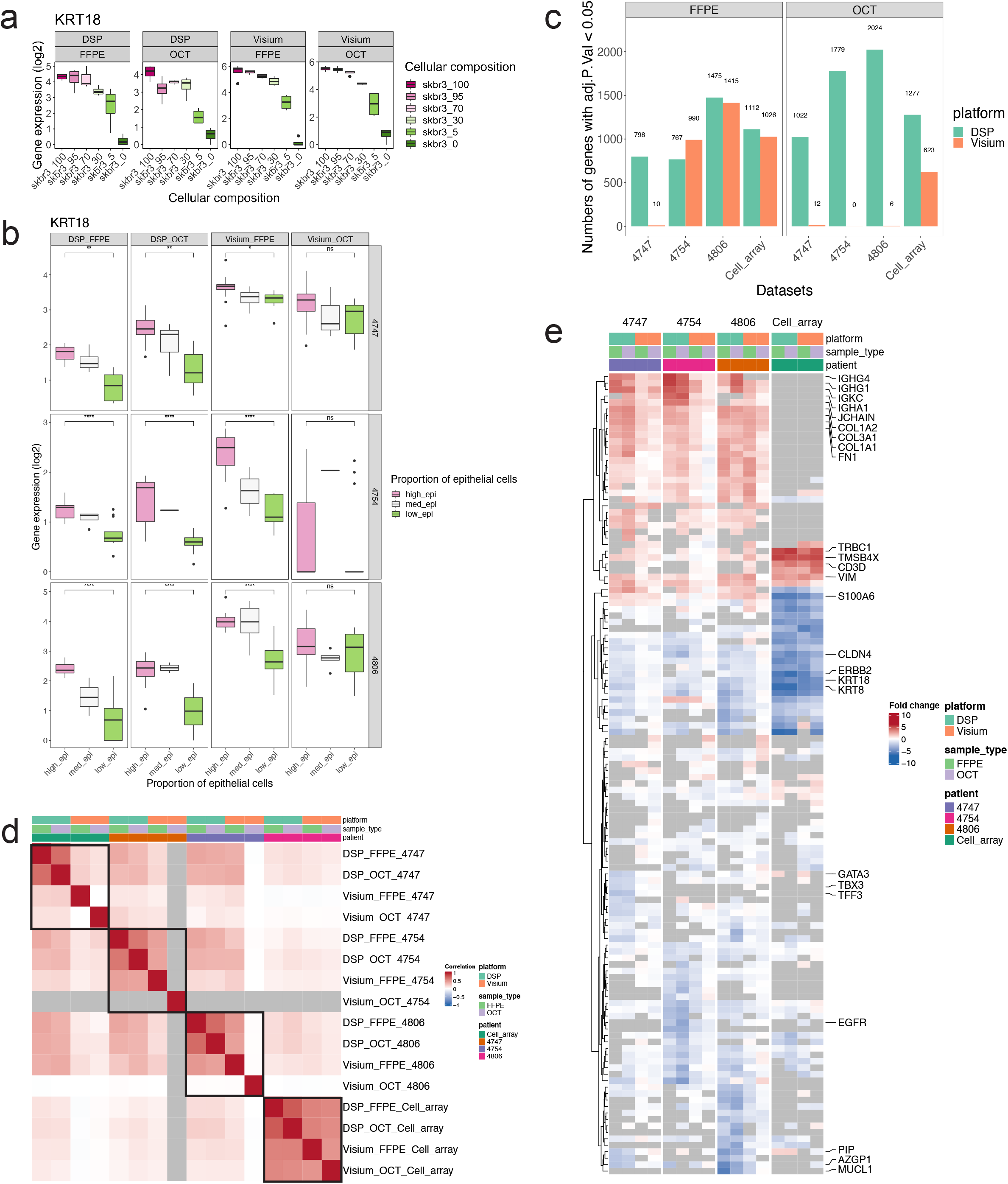
Comparison of the DSP and Visium platforms in detecting gene expression changes. Visium mimic AOIs in DSP data and Visium spots from the matching region were used for the analyses. All gene expression data was down sampled to manufacturer’s recommendation at a per sample level. (**a**) Normalised expression of cell markers in the cell array samples detected by the DSP or Visium assays. ns: non-significant. *p<0.05, **p<0.01, ****p<0.0001, student t-test. (**b**) Normalised expression of cell markers in tissue samples detected by the DSP or Visium assays. (**c**) The number of DEGs detected by the DSP and Visium assays in each sample. A gene is considered to be differentially expressed if adjusted p < 0.05. (c) Correlation of fold change of DEGs detected by the DSP or Visium platforms in each sample. (**e**) Fold changes of top 10 DEGs detected by the DSP and Visium platforms in each sample. DEGs considered to be enriched in the non-epithelial AOIs / spots were given positive fold changes while DEGs enriched in the epithelial AOIs / spots have negative fold changes.

To investigate why the Visium data did not markedly reflect gene expression changes between tissue compartments with different cellular composition, we evaluated the expression of cell lineage specific marker genes in individual Visium spots in sample 4747 (FFPE). This showed that the expression of these genes are not completely restricted to the corresponding pathology annotated tissue regions (Fig. S9). This is likely due to infiltration of the tumour by immune / stromal cell types (i.e. PTPRC and COL1A1 expression). However, some expression of epithelial markers such as KRT8 and KRT18 was also detected in spots annotated as stroma, despite the limited presence of cancer cells in these spots as revealed by the H&E image (Fig. S9). While the exact cause of such observation is still unclear, a recent study has suggested that transcripts or probes in Visium assays might diffuse into adjacent spots during tissue permeabilization leading to an effect termed as spot swapping [29]. However, the extent to which this influences the current Visium datasets remains uncertain. For the Visium OCT data, the overall low detection of DEGs across tissue samples could be due to the uneven distribution of UMIs across the genes as observed in Fig. 2c-e.

To more quantitively compare the performance of DSP and Visium in detecting the difference in gene expression between different regions, we conducted differential gene expression (DE) analysis between the high epithelial and low epithelial AOIs/spots collected by each assay. For the cell array samples, only data collected from 100% SKBR3 and 100% Jurkat cells was used. For tissue samples, AOIs/spots were manually annotated based on tissue morphology. We first compared the numbers of differentially expressed genes (DEGs) detected by each assay, with an adjusted p-value less than 0.05. In general, the DSP platform generates fairly similar results across all samples tested (Fig. 3c). The number of DE genes were comparable between the DSP and Visium FFPE solutions for sample 4754, 4806 and the cell array. However, few DEGs were detected with the Visium FFPE solution for sample 4747 and all tissue samples processed using the OCT Visium assay (Fig. 3c).

We then tested the concordance between DSP and Visium DE results by computing the Pearson’s correlation score between the fold changes of significant DE genes. This showed high correlation between all platforms on the cell-array samples (Fig. 3d). We also observed good correlation between DSP data from matching FFPE and OCT samples in all tissue samples tested. The correlation of FFPE Visium results with DSP data was also good in samples 4754 and 4806 (above 0.5) but poor in 4747. OCT Visium had poor correlation with all other datasets in tissue samples (Fig. 3d).

In addition to the overall pattern, we also examined the biology revealed by the DE analysis. The fold change of the top 10 DEGs identified by each assay in each sample was plotted as a heatmap. For data generated from samples FFPE 4747, OCT 4747, OCT 4754 and OCT 4806 by the Visium platform, the fold changes detected appear to be smaller than the fold changes detected by other assays on the same samples (Fig. 3e). This is in line with the previous analysis results in which limited numbers of DEGs were confidently detected by the Visium assays in these samples (Fig. 3c). Nonetheless, the overall fold change pattern is consistent across all datasets. In the cell-array dataset we observed strong DE of markers related to Jurkat (CD3D, TRBC1, TMSB4X) or SKBR3 (ERBB2, KRT8, KRT18) cells (Fig. 3e). Epithelial-depleted tissue regions featured genes encoding immunoglobulin and collagen genes, consistent with enrichment of fibroblasts and B cells in those regions, however many genes enriched in SKBR3 were also found to be enriched in regions with high epithelial content, reflecting the epithelial nature of SKBR3 breast cancer cells (Fig. 3e). We also identified sample-specific gene clusters. For example, we observed enrichment of GATA3 and TFF3 in the cancer regions of 4747, consistent with its clinical classification as a luminal breast cancer.

### Comparison of the DSP and Visium platforms on resolving fine tissue structures

So far, we have focused on unsegmented DSP AOIs when making direct comparisons between the DSP and Visium assays. However, a unique feature of the DSP platform is its ability to collect transcriptomic profiles of different cell types separately based on fluorescence masking. We tested this ability of the DSP platform using segmented AOIs targeting epithelial or non-epithelial segments based on the staining of anti-pan-cytokeratin antibody. In comparison to the gene expression data collected using DSP segmentation, Visium spots located in regions with matching morphology were manually selected and separated into the epithelial and non-epithelial group based on cellular composition (Fig. 1c iii-iv).

We first evaluated the purity of DSP segmentation using cell array samples. As shown previously (Fig. 3a), we observed a good concordance between the expression of cell markers and the proportion of SKBR3 and Jurkat cells when using Visium given the expression of both cell lines were captured together using the Visium assays (Fig. S10). However, on the other hand, we observed enrichment of cell markers in the corresponding DSP segments irrespective of the mixing proportion of Jurkat and SKBR3 cells confirming the enrichment of cell type-specific transcriptomic profile through DSP segmentation (Fig. S10).

To evaluate the purity of DSP segmentation, directly from the whole transcriptomic profile, rather than relying on a handful of cell type markers, we used deconvolution to infer the proportion of Jurkat and SKBR3 cells. Encouragingly, we observed a similar pattern in deconvolution results, where DSP segments in FFPE cell array samples were predicted to contain almost exclusively Jurkat or SKBR3 cells in the corresponding segments, whereas the proportion detected using the Visium assay changes as the mixing proportion changes between Jurkat and SKBR3 cells (Fig. 4a). However, we did notice that the separation is not as clear in data collected using the DSP OCT assay. As the proportion of SKBR3 and Jurkat changes, the predicted cell proportion changed correspondingly (Fig. 4a).

**Figure 4:**
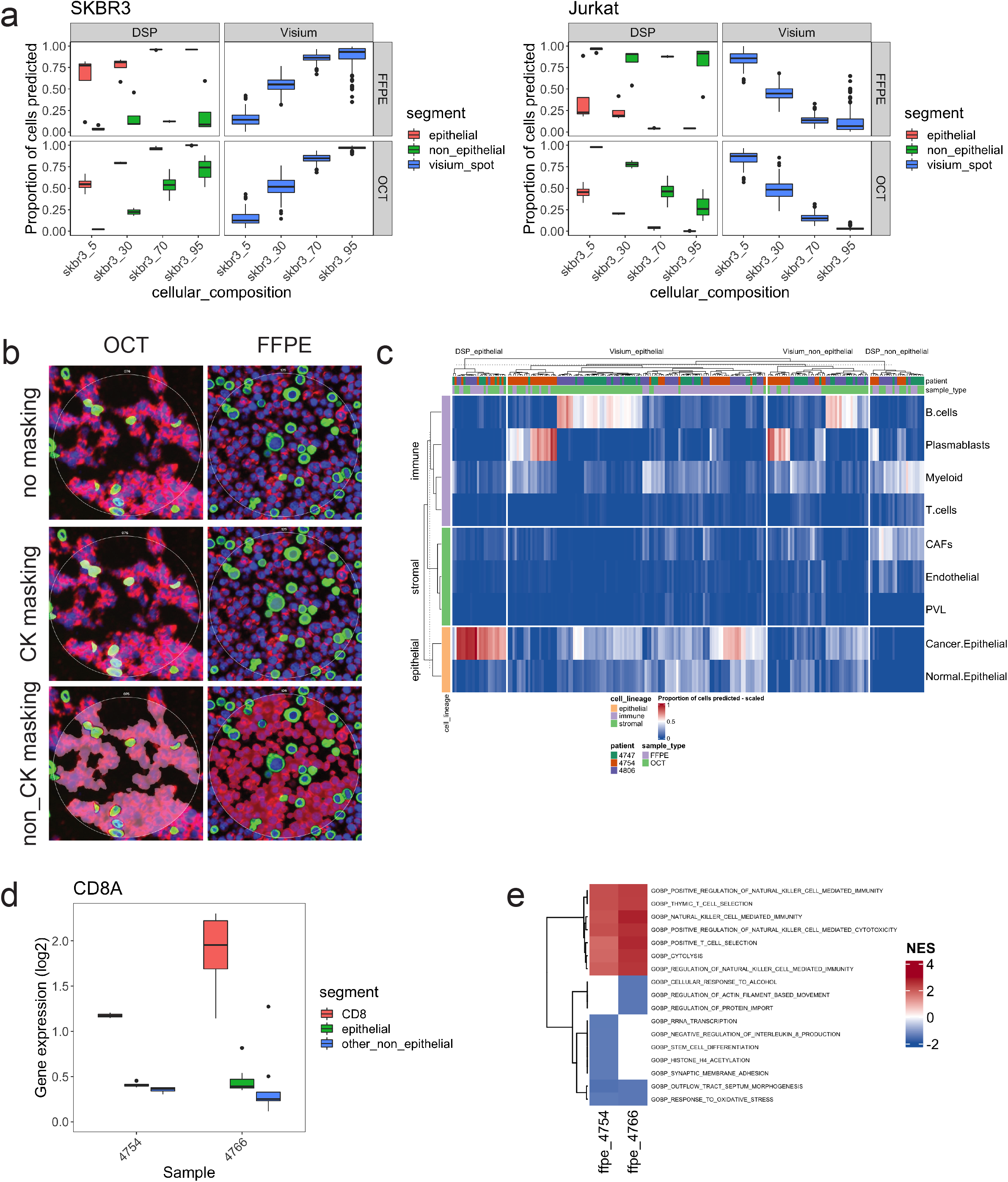
Segmentation by the DSP platform allow profiling of more specific gene expression features. (**a**) Predicted proportion of SKBR3 and Jurkat cells in segmented DSP AOIs and Visium spots in cell array samples. (**b**) Example of segmented DSP AOIs on cell array samples. All cells were labelled with SYTO13 for nuclei stain. Jurkat and SKBR3 cells were labelled with anti-CD45 (red) or pan-cytokeratin (green) antibodies respectively. Epithelial or non-epithelial DSP AOIs were then sampled based on the fluorescent signals in a sequential manner. (**c**) Predicted proportion of cancer and TME cell types in segmented DSP AOIs and Visium spots in tissue samples. (**d**) Normalised expression of CD8A in CD8 and non CD8 segments in DSP data. (**e**) NES of top 5 significantly dysregulated GOBP pathways identified between CD8 segments and adjacent non-CD8 TME segments by the DSP platform. Significance threshold was set as q < 0.25.

The results above seem to indicate that the segmentation works better in FFPE samples than in the OCT samples for the DSP platform. To better understand the cause of such observations, we evaluated the immunofluorescence images to understand the variations in the FFPE and OCT cell array samples. Of note, cells in the FFPE cell array appear to be forming a relatively uniformed single layer, while cells in the OCT cell array seem to have aggregated into strips (Fig. 4b). Given that segmentation was only conducted in two dimensions, it is possible that there are other cell types above or below the targeted cell type, causing contamination of the gene expression signal and less clear separation using the IF-based segmentation.

We next extended our comparisons to breast cancer tissue samples. In line with the results above, we observed enrichment of marker gene expression in corresponding tumour or non-tumour DSP segments (Fig. S11). In many samples, a difference can also be observed between Visium spots annotated as epithelial and non-epithelial (Fig. S11). We also predicted the cellular composition in our spatial datasets using gene expression signatures defined in our published breast cancer single cell RNA-Seq dataset [5]. In both OCT and FFPE DSP data, the tumour segments were predicted to contain almost exclusively epithelial cancer cells, which were absent in the non-tumour segments (Fig. 4c), whereas Visium spots annotated as epithelial or non-epithelial are predicted to contain immune and stromal cell types, along with epithelial cancer signatures (Fig. 4c), as would be expected in a tumour.

In the analyses above, we profiled regions with well compartmentalised tissue structures and a clear tumour-stroma interface. We then challenged the DSP platform by targeting more specific cell types, namely CD8 T cells, in the tumour microenvironment of two samples. We focused on biological AOIs as shown in Fig. 1d. The transcriptomic profiles of CD8 T cells were collected through segmentation based on immunofluorescence signal of an anti-CD8 antibody. The transcriptomic profiles of adjacent tumour cells and non-CD8 TME cell types were also collected.

To test the purity of segmentation, we examined the expression of cell markers, including CD8A. From this, we observed enriched CD8A gene expression in CD8 segments as compared to the epithelial segments or non-epithelial-non-CD8 segments collected in the same region (Fig. 4d). We also conducted DE analysis between CD8 segments and adjacent non-CD8 TME AOIs, which showed significant enrichment of T cell-related pathway activity (Fig. 4e). However, the expression of myeloid cell markers CD14 and CD68 as well as B cell marker JCHAIN were also high in the CD8 segment, at a level comparable to adjacent non-CD8 TME segments (Fig. S12). While the exact cause of the contamination in the CD8 transcriptomic profile is unclear, it may be caused by interactions between immune and stromal cell types that cannot be fractionated through segmentation, highlighting a challenge for DSP segmentation in obtaining pure transcriptomic profiles in complex tissues.

### Comparison of the DSP and Visium platforms in profiling the molecular landscape of tumours

Previous analyses were mainly focused on certain regions of the tumour partially due to the nature of the DSP platform which allows the deeper profiling of specific areas with rich morphological features. In contrast, the Visium platform requires minimal guidance on area selection and allows non-biased characterisation of the tissue at relatively high spatial resolution. This potentially provides a data-driven, hypothesis-generating approach to characterising the molecular landscape of tissue samples. We examined this feature of the Visium platform to investigate the spatial heterogeneity of cancer cells in our samples. We selected Visium spots with high tumour content through pathological evaluation and inferred the cancer cell composition in these spots using single-cell RNA-Seq transcriptomic signatures [5] (Fig. 5a). The luminal A/luminal B, HER2E and basal subtypes defined through single-cell analysis generally correlates with ER+, HER2+ and TNBC breast cancers in the clinical setting. A proliferating/cycling cancer signature was also defined to reflect the active proliferating cell state of breast cancer cells in the single-cell dataset [5]. For direct DSP and Visium comparison, we predicted the cancer cell proportions in DSP AOIs and Visium spots from matching regions on each sample. In addition, results from all Visium spots were included to evaluate the global pattern of cancer composition across each sample. We observed a good concordance between the predicted molecular subtypes and the known clinical subtypes for 4747 (ER+) and 4754 (TNBC) (Fig. 5a) whereas, 4806 (TNBC) was predicted to mainly contain cancer cells of the HER2E subtype when using both DSP and Visium platforms (Fig. 5a). This is not surprising as discordance between clinical and molecular subtype is observed in up to 38% of breast cancer cases [30]

**Figure 5:**
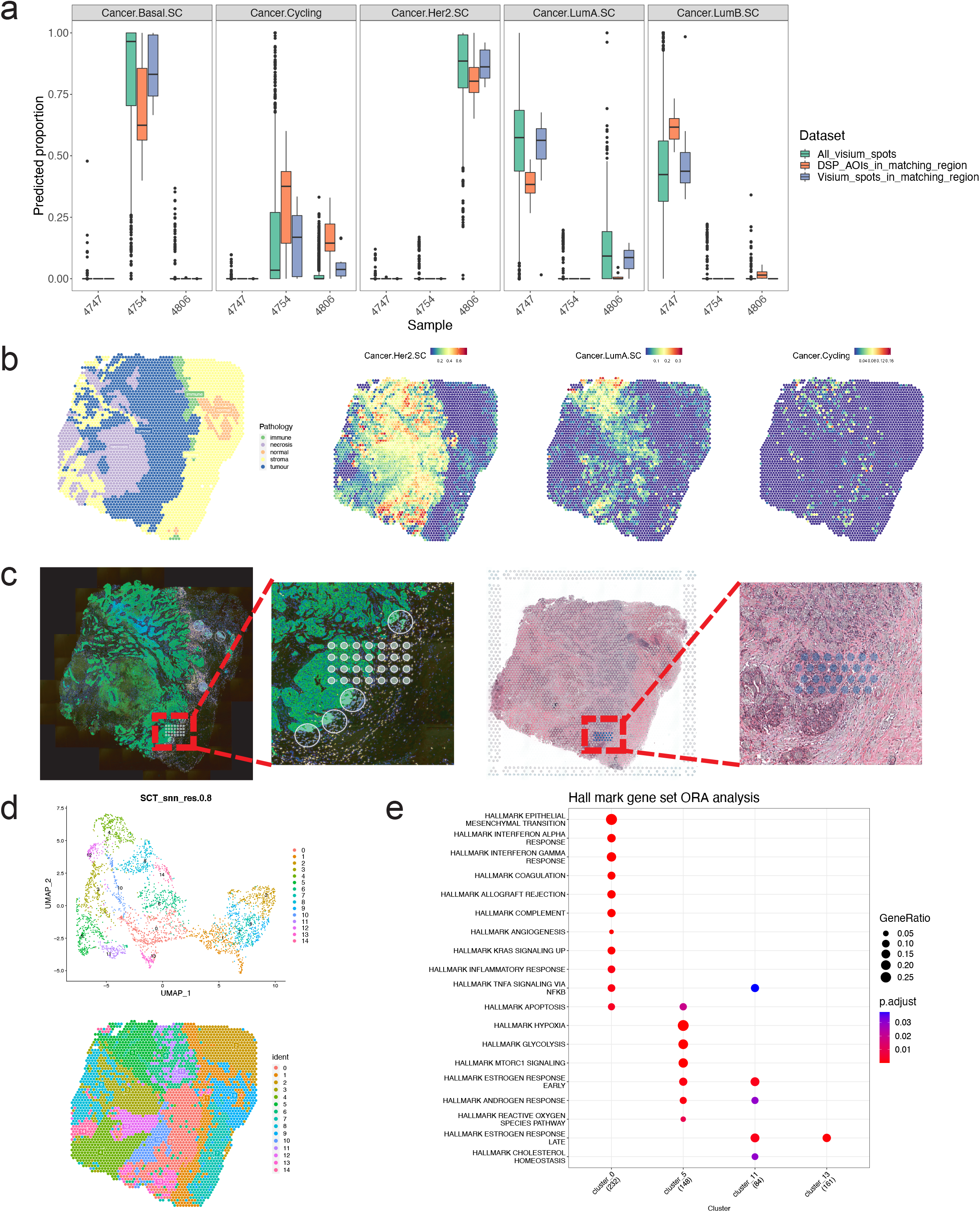
The Visium platform generates a transcriptomic map facilitating unbiased heterogeneity exploration. (**a**) Predicted proportion of breast cancer subtypes DSP and Visium data. All DSP and Visium AOIs /spots with the same size were used in the analysis. To account for spatial heterogeneity in the samples, Visium spots from regions with matching DSP AOIs were annotated and plotted as the 3^rd^ group. Only AOIs / spots with high cancer proportion by pathology were included. (**b**) pathology annotation of FFPE 4806 in Visium data and the spatial distribution pattern of predicted breast cancer subtypes by deconvolution in this sample. (**c**) Illustrative images of the location of DSP AOIs and Visium spots in FFPE 4806. (**d**) Clustering of Visium spots based on gene expression profiles and spatial projection of the clustering results. (**e**) ORA analysis between clusters with high cancer cell proportion in Visium data.

To validate the cell type deconvolution prediction, we examined the expression of common breast cancer subtype markers [5,31] and observed high expression of luminal cancer markers ESR1 and TFF1 in the cancer region of sample 4747 and basal cancer markers KRT6B and EGFR in sample 4754 (Fig. S13). For sample 4806, we did observed expression of HER2 cancer markers such as ERBB2 and GRB7 though the expression is not outstanding when comparing to the other two samples (Fig. S13). However, importantly, only minimal expression of luminal and basal cancer markers was observed in 4806 (KRT6B expression in 4806 was mainly associated with tissue necrosis) (Fig. S13) suggesting that this sample should indeed be classified as a breast cancer of HER2 molecular subtype.

While the two platforms were broadly concordant, we observed some differences in prediction between the DSP and Visium assays. In addition to HER2 breast cancer cells, the Visium platform also predicted sample 4806 to contain cancer cells of luminal A subtype (Fig. 5a). Indeed, we observed some correlation in spatial distribution of several luminal cancer markers including KRT8, KRT18 and TFF3 with the predicted luminal A signatures (Fig. S14). Interestingly, the spatial distribution of luminal A cancer cells was more heterogeneous (Fig. 5b) than Her2E cells, with the signature enriched in regions at the top of the tissue. This area was not sampled by the DSP AOIs, which were in a distant region of this tissue (Fig. 5c), highlighting the strength of the Visium platform to enable more comprehensive, practical, exploratory sampling of wider tumour regions. However, in regions covered by both the DSP and Visium data, both platforms demonstrate high concordance in resolving the molecular profiles of tumour cells.

To better understand the biological nature of the tumour cells in 4806, we then clustered the Visium spots, in an unsupervised manner, using the spatial gene expression profiles. In total, 15 clusters were identified from all Visium spots on FFPE 4806 (Fig. 5d). Among these clusters, C0, C3, C5, C10, C11, C12 and C13 were predicted, by deconvolution, to be comprised of over 50% epithelial cells (Fig. S15). Of these, C3, C10 and C12 were found to be located in regions affected by necrosis and therefore excluded from the downstream analysis. For the remaining 4 clusters (C0, C5, C11 and C13), C0 is located at the edge of the tumour mass, adjacent to a clustered region with high immune cell composition (C1) (Fig. 5b,d; Fig. S16), C5 is located in the region predicted to contain the highest luminal A signature (Fig. 5b,d; Fig. S16). The remaining 2 clusters C11 and C13 are located in regions with mainly HER2 cancer signatures (Fig. 5b,d; Fig. S16).

We then characterised the biological processes enriched in each cancer cluster, computing the top differentially expressed genes in each cluster. The top 10 (if available) significantly dysregulated pathways in each cluster were then selected, which showed a large enrichment of immune related pathways in C0, compared to the others (Fig. 5e). The spatial proximity of this cluster, to C1, which has high predicted immune cell composition (Fig. S15) suggests molecular interactions between tumour and adjacent immune cells. We also observed enrichment of estrogen related signalling in clusters C5, C11 and C13 which is in line with the predicted presence of Luminal A cancer cells in these spatial locations (Fig. 5e). Interestingly, we noticed that both the androgen response and apoptosis gene set activities were significantly upregulated in C5. Previously literature has suggested that AR activity may have a tumour-suppressive function in ER positive breast cancer cells [32]. While the exact molecular mechanism driving AR activation in C5 remains to be further evaluated, our results have demonstrated the capacity of the Visium platform in performing non-biased, data driven characterisation of tissue heterogeneity.

## Discussion

The DSP and Visium technologies, along with other platforms such as Slide-seq [33], MERFISH [34] and Seq-FISH [35], are driving a revolution in our ability to spatially profile biology at whole transcriptome molecular resolution. Both the DSP and Visium platforms have sophisticated designs and are leading platforms in the spatial analysis of heterogeneity in tissue [4,36,37,38]. However, a direct evaluation of the performance of DSP and Visium platform is still missing, making platform selection a difficult task for researchers entering this area. In this study, we utilised a collection of well-controlled cell line and tissue samples to address this gap and provide a better understanding of the strengths and limitations of these platforms for spatial transcriptomics and oncology research.

Direct comparison of the DSP and Visium platforms was conducted using AOIs/spots of equivalent size and number. We observed a high correlation in the cell array samples where the cellular composition was precisely controlled and the sample structure was relatively simple, but discordance between DSP and Visium was seen when profiling breast cancer tissues. Most surprising was the discordance when using Visium on OCT processed samples, where the level of gene detection as well as the difference in gene expression between distinct cellular regions was significantly lower. The reason for these discrepancies are unclear as the QC parameters (such as reads per spot and genes per spot) of the OCT Visium datasets are within the expected ranges and established experimental protocols were followed. While tissue permeabilisation does not appear to be the cause in this study, effectively evaluating and balancing the strength of the RNA footprint obtained by imaging is certainly a challenging step in the OCT Visium workflow. In addition, this is recommended to be performed on a per-sample basis, which increases experiment cost, time, tissue required, and reduced the throughput of the OCT Visium workflow. Despite these challenges this is the only platform, among those compared, that does not require the use of targeted RNA probes, thereby enabling capture of all intrinsic RNA molecules with poly-A sequences. This can be particularly valuable for profiling transcripts whose nucleotide sequences are variable, for example, TCR or BCR [39].

Unlike the OCT Visium datasets, the FFPE Visium datasets detected large numbers of genes within each spot. In addition, investigation of immunoglobulin gene expression in the cell array samples revealed essentially no background in the FFPE Visium assay. In contrast, this experiment did identify non-specific binding of probes in the DSP assays. Genes with non-specific detection can potentially be filtered out using the built-in non-targeting control probes. However, this may also impact true signal with relative weak intensity. More sophisticated background removal methods have been proposed [24,40], however, the performance of these algorithms remain to be further tested. It is worth noting that while Visium samples were sequenced extensively (achieving more than twice the recommended sequencing depth), the saturation of the gene expression library was only around 40%, indicating the possibility to further improve gene detection with deeper sequencing.

The FFPE and OCT DSP data demonstrated high consistency in gene detection across all samples. OCT DSP data seems to perform better than FFPE DSP data with more genes detected per AOI and more DEGs detected between high epithelial and low epithelial regions. Given that OCT samples generally have better RNA quality than FFPE samples, this is probably as expected and a reflection of the variations in tissue quality between assays.

In addition to a controlled, direct technical comparison between DSP and Visium, we also explored the unique strengths of each platform. For instance, the DSP platform allows the separation of transcriptomic profiles of closely located cell populations using morphology masking. However, the results seem to be more promising in well compartmentalised tissue structures (such as tumour versus non-tumour) compared to regions where the boundaries between different cell populations are less clear (such as between tightly interacting CD8 T cells and myeloid cells). In contrast, the Visium platform averages expression of closely-interacting cells.

However, the Visium platform provides good coverage at relatively high resolution across the whole sample in the capture area, making it more suited to unbiased profiling of tumour heterogeneity across larger tissue areas. We suggest these observations highlight the scenarios where the unique strengths of the DSP and Visium assays should be applied.

While not the focus of this study, the DSP and Visium platforms also vary in several other features. Firstly, the morphology masking antibodies used in the DSP workflow typically require optimisation prior to the experiment. For example, in the current study, a CD8 antibody was conjugated with fluorescent dye with concentration titrated to obtain the optimal image for DSP experiments. In addition, a different CD45 antibody was used to label Jurkat cells in the cell array samples due to cross reactivity of the default CD45 antibody from the DSP morphology marker kit with SKBR3 cells.

Another variation between the platforms is related to sequencing library construction. For Visium, samples from different spatial spots are pooled together prior to library amplification, allowing for easier handling of the samples. These samples were then amplified together, within the same PCR reaction, minimising batch effects. In contrast, DSP libraries require more labour-intensive handling of samples stored in 96-well plates. Several plates are required for large experiments such as the current study, which may lead to bias or human error (such as pipetting error) when processing individual samples separately.

Thirdly, the capture areas of the DSP and Visium platforms have different dimensions. Samples in the current study were intentionally bio-banked to fit the capture area on Visium slides (6.5mm x 6.5mm). However, common histological FFPE blocks can reach 2cm x 2cm in size if not bigger or are in specific shapes such as biopsy samples which are 1-2mm in diameter but 1-2cm in length. It is therefore impossible to fit all parts of the samples into the capture area on Visium slides. Additional trimming or handling is required for these samples, increasing the labour-cost of Visium experiments. On the other hand, the capture area in the DSP platform is substantially larger (36.2mm x 14.6mm), making it more compatible for this type of tissue and potentially for TMA samples.

There are some additional caveats of our study. Firstly, the FFPE Visium data was generated by the manufacturer who developed the technology. Therefore, the high data quality of FFPE Visium data in the current study could represent over-optimised conditions that are challenging to replicate in a typical laboratory. Secondly, due to unknown reasons, the OCT Visium data appears suboptimal meaning that the comparisons to these datasets maybe considered less conclusive. Thirdly, the DSP AOIs studied in this analysis were mainly located in a confined region of the tissue samples. While this does reflect a common workflow for the DSP platform, in requiring prior knowledge of the sample to be studied, the low coverage of DSP AOIs across the tissue samples limited our ability to systematically compare the two platforms’ ability to detect regional differences in tumour heterogeneity. Finally, the sample size used in this study is relatively small. While the 3 breast cancer tissue samples do cover luminal, HER2 and TNBC subtypes of breast cancer, more samples across more diverse tissue and cancer types will be required to exhaustively assess whether the current observations are maintained.

## Conclusion

In this study, we performed controlled comparisons between the DSP and Visium platforms to assess their ability to capture spatially resolved transcriptomic features in breast cancers. We show that the two platform generate broadly comparable results using carefully controlled conditions and samples. We propose that the Visium platform is more suitable in generating a non-biased transcriptomic landscape of the whole tissue. This enables the identification of cell populations harbouring unique gene expression signatures but with seemingly similar morphological features to other cells. To complement this, the DSP platforms prevails in deep molecular profiling of known regions with prior knowledge of tissue regions of interest and is more suited to addressing hypothesese. Clearly there are advantages to combining DSP and Visium assays in the same study, starting with discovery and hypothesis generation using the Visium platform and followed by hypotheses testing or validation using the DSP assays.

It’s also worth noting that neither DSP nor Visium provides spatially resolution at the single-cell level. To bridge this gap, new platforms based on optical imaging or high-density spots or arrays of beads are in development or being commercialised [33,41,42]. While these technologies promise improved spatial resolution they are still mostly limited in transcriptomic coverage when compared to the DSP and the Visium platform. Until a technology is developed that can deliver the trifecta of wide transcriptomic coverage, single cell resolution and large capture areas the DSP and Visium platforms look set to remain as two key technologies for generating spatial whole transcriptomic profiles, furthering our knowledge of the spatial molecular nature of the tissue samples and fuelling the advancement of research, treatment and care in various disease settings.

## Figure legends

**Supplementary figure 1: DSP AOI size and gene detection.** Only data from DSP size gradation AOIs were used in the analysis. Boxplot showing changes in the numbers of genes detected per AOI as the AOI size changes.

**Supplementary figure 2: QC of data from Visium mimic AOIs (DSP) or Visium spots (Visium).** (**a-c**) QC of original DSP and Visium datasets. (**a**) Average number of raw reads per μm^2^ collected by the DSP assays in each sample before down sampling. (**b**) Average number of raw reads per spot collected by the Visium assays in each sample before down sampling. (**c**) Average sequencing saturation of DSP and Visium data before down sampling. (**d-f**) QC of down sampled DSP and Visium datasets. (**d**) Average number of raw reads per μm^2^ collected by the DSP assays in each sample after down sampling. (**e**) Average number of raw reads per spot collected by the Visium assays in each sample after down sampling. (**f**) Average sequencing saturation of DSP and Visium data after down sampling.

**Supplementary figure 3: Distribution of UMI across all genes detected in each dataset.** Only DSP visium-mimic AOIs and matching visium spots were used for the plot. All genes with UMI detected in each sample were ranked based on the total numbers of UMI detected for each gene and plotted on the x axis. The cumulative proportion of all UMI collected in each sample was plotted on the y axis. Top 5 genes with the most UMI per gene collected were annotated. The numbers of genes contributing to 50% and 75% of all UMIs collected were also labelled.

**Supplementary figure 4: non-specific detection.** Only data from Visium mimic AOIs in DSP data from cell array samples and Visium spots in the matching region was used in this analysis. (**a-c**) heatmap of counts of immunoglobulin heavy chain genes and marker genes detected by the DSP and Visium assays. (**a**) raw counts from both the DSP and Visium assay were plotted. (**b**) DSP counts were filtered by geometric mean of non-targeting control probes. Raw counts were plotted for Visium data. (**c**) DSP counts were filtered by limit of quantitation. The same raw count were plotted for Visium data.

**Supplementary figure 5: Sensitivity of the DSP platform with background filtered counts.** (**a**) Numbers of genes detected per AOI / spot. DSP data was filtered using the geometric mean of all non-targeting probe readings. Any gene with count above 0 after filtering was considered detected. Raw Visium data was plotted. (**b**) DSP matrix sparsity using background filtered counts. (**c**) UMI distribution plots of DSP data using background subtracted counts.

**Supplementary figure 6: Sensitivity of the DSP platform with LOQ filtered counts.** (**a**) Numbers of genes detected per AOI / spot. DSP data was filtered using the LOQ threshold. Any gene with count above 0 after filtering was considered detected. Raw Visium data was plotted. (**b**) DSP matrix sparsity using LOQ filtered counts. (**c**) UMI distribution plots of DSP data using LOQ subtracted counts.

**Supplementary figure 7: Normalised expression of cell markers in Visium mimic AOIs and matching Visium spots from cell array samples.**

**Supplementary figure 8: Normalised expression of cell markers in Visium mimic AOIs and matching Visium spots from tissue samples.** ns: non-significant. *p<0.05, **p<0.01, ***p<0.001, ****p<0.0001, student t-test.

**Supplementary figure 9: pathology annotation and spatial expression patterns of marker genes in FFPE 4747 profiled by the Visium platform.** (**a**) Visium spots used for direct DSP and Visium comparison. (**b**) pathology annotation of the tissue sample. (**c**) Normalised expression of cell markers.

**Supplementary figure 10: Normalised expression of cell markers detected using segmented DSP AOIs or Visium spots in the matching region in cell array samples.**

**Supplementary figure 11: Normalised expression of cell markers detected using segmented DSP AOIs or Visium spots in the matching region in tissue samples.**

**Supplementary figure 12: Normalised expression of cell markers in CD8 segments and adjacent epithelial and non-CD8 TME segments in DSP data.**

**Supplementary figure 13: Normalised expression of common breast cancer subtype markers in FFPE Visium data.**

**Supplementary figure 14: Normalised expression of luminal breast cancer markers in Visium data from FFPE 4806.**

**Supplementary figure 15: Proportion of major cell types predicted in each Visium cluster from FFPE 4806.**

**Supplementary figure 16: Proportion of breast cancer subtypes predicted in cancer clusters in Visium data from FFPE 4806.**

## List of abbreviations

AOI: area of illumination
BCR: B cell receptor
DCC: digital count conversion
DE: differential gene expression
DEG: differentially expressed gene
DSP: digital spatial profiling
FFPE: formalin-fixed paraffin embedded
HER2E: HER2 enriched
LOQ: limit of quantification
NBF: neutral-buffered formalin
OCT: optimal cutting temperature compound
ORA: over-representation analysis
QC: quality control
ROI: region of interest
TCR: T cell receptor
TME: tumour microenvironment
TNBC: triple negative breast cancer
UMAP: uniform manifold approximation and projection
UMI: unique molecular identifier

## Declarations

### Ethics approval and consent to participate

The human breast cancer samples used in this study were collected following protocols x13-0133 and x19-0496. Ethical approval for this study was acquired by the Sydney Local Health Districts Ethics committee - Royal Prince Alfred Hospital zone. Consent for the use of tissue samples was obtained from all patients prior to collection, and both tissues and data were de-identified as per approved protocol.

### Consent for publication

Not applicable

### Availability of data and materials

The datasets supporting the conclusions of this article are being uploaded to the European Genome-phenome Archive (EGA) repository. The code and scripts used to generate results of this article are being uploaded to a github repo (https://github.com/Swarbricklab-code/NvV_paper_code_2023).

All datasets and codes are currently being uploaded and are available upon reviewer request and will be made publicly available prior to publication.

### Authors’ contributions

A.S. conceived the project and directed the study. T.W. and A.S. wrote the manuscript. All authors reviewed the drafting of the manuscript. E.L., S.O.T. and K.H organised the access to breast cancer patient tissue. K.H. collected clinical samples. K.H. and J.Y. prepared the cell line samples. T.W and K.H conducted the Nanostring DSP experiments. C.C and D.K conducted Visium OCT experiment. S.O., T.W. and K.H. conducted pathology annotation of the tissue samples. T.W. and J.R. performed and interpreted the analysis of Nanostring and Visium data. D.R. and N.B. supervised the data analysis. G.A. and J.P. provided intellectual input. The authors read and approved the final manuscript.

## Acknowledgements

We thank Anaiis Zaratzian and Andrew Da Silva from the histology facility in the Garvan Institute of Medical Research for their support on preparing FFPE tissue sections for the FFPE DSP assay.

We thank the Garvan Molecular Genetics team for their support on conducting RNA integrity assessment for OCT tissue samples.

We thank Jason Reeves, Jingjing Gong, Marshall Feterl, Swati Ranade, Nicholas Confuorto and Kit Fuhrman from Nanostring for their advice on conducting DSP experiments and analysing DSP data.

We thank the 10X Genomics R&D team for generating Visium FFPE data. We also thank Stephen Williams and Sarah Taylor from 10X Genomics for their advice on analysing Visium data.

## References

1. Vitale I, Shema E, Loi S, Galluzzi L. Intratumoral heterogeneity in cancer progression and response to immunotherapy. Nat Med. 2021;27:212–224.

2. Moses L, Pachter L. Museum of spatial transcriptomics. Nat Methods. 2022;19:534–546.

3. Bergholtz H, Carter JM, Cesano A, Cheang MCU, Church SE, Divakar P, Fuhrman CA, Goel S, Gong J, Guerriero JL, et al. Best Practices for Spatial Profiling for Breast Cancer Research with the GeoMx Digital Spatial Profiler. Cancers (Basel). 2021;13.

4. Hwang WL, Jagadeesh KA, Guo JA, Hoffman HI, Yadollahpour P, Reeves JW, Mohan R, Drokhlyansky E, Van Wittenberghe N, Ashenberg O, et al. Single-nucleus and spatial transcriptome profiling of pancreatic cancer identifies multicellular dynamics associated with neoadjuvant treatment. Nat Genet. 2022;54:1178–1191.

5. Wu SZ, Al-Eryani G, Roden DL, Junankar S, Harvey K, Andersson A, Thennavan A, Wang C, Torpy JR, Bartonicek N, et al. A single-cell and spatially resolved atlas of human breast cancers. Nat Genet. 2021;53:1334–1347.

6. Bassiouni R, Gibbs LD, Craig DW, Carpten JD, McEachron TA. Applicability of spatial transcriptional profiling to cancer research. Mol Cell. 2021;81:1631–1639.

7. Asp M, Bergenstrahle J, Lundeberg J. Spatially Resolved Transcriptomes-Next Generation Tools for Tissue Exploration. Bioessays. 2020;42:e1900221.

8. Lewis SM, Asselin-Labat ML, Nguyen Q, Berthelet J, Tan X, Wimmer VC, Merino D, Rogers KL, Naik SH. Spatial omics and multiplexed imaging to explore cancer biology. Nat Methods. 2021;18:997–1012.

9. Hao Y, Hao S, Andersen-Nissen E, Mauck WM, 3rd, Zheng S, Butler A, Lee MJ, Wilk AJ, Darby C, Zager M, et al. Integrated analysis of multimodal single-cell data. Cell. 2021;184:3573–3587 e3529.

10. Nicole Ortogero, Zhi Yang, Ronalyn Vitancol, Maddy Griswold, Henderson D. GeomxTools: NanoString GeoMx Tools. 2022.

11. Seqtk: a fast and lightweight tool for processing FASTA or FASTQ sequences [https://github.com/lh3/seqtk]

12. Ritchie ME, Phipson B, Wu D, Hu Y, Law CW, Shi W, Smyth GK. limma powers differential expression analyses for RNA-sequencing and microarray studies. Nucleic Acids Res. 2015;43:e47.

13. Gu Z, Eils R, Schlesner M. Complex heatmaps reveal patterns and correlations in multidimensional genomic data. Bioinformatics. 2016;32:2847–2849.

14. Andersson A, Bergenstrahle J, Asp M, Bergenstrahle L, Jurek A, Fernandez Navarro J, Lundeberg J. Single-cell and spatial transcriptomics enables probabilistic inference of cell type topography. Commun Biol. 2020;3:565.

15. Wolf FA, Angerer P, Theis FJ. SCANPY: large-scale single-cell gene expression data analysis. Genome Biol. 2018;19:15.

16. Satija R, Farrell JA, Gennert D, Schier AF, Regev A. Spatial reconstruction of single-cell gene expression data. Nat Biotechnol. 2015;33:495–502.

17. Danaher P, Kim Y, Nelson B, Griswold M, Yang Z, Piazza E, Beechem JM. Advances in mixed cell deconvolution enable quantification of cell types in spatial transcriptomic data. Nat Commun. 2022;13:385.

18. Mootha VK, Lindgren CM, Eriksson KF, Subramanian A, Sihag S, Lehar J, Puigserver P, Carlsson E, Ridderstrale M, Laurila E, et al. PGC-1alpha-responsive genes involved in oxidative phosphorylation are coordinately downregulated in human diabetes. Nat Genet. 2003;34:267–273.

19. Subramanian A, Tamayo P, Mootha VK, Mukherjee S, Ebert BL, Gillette MA, Paulovich A, Pomeroy SL, Golub TR, Lander ES, Mesirov JP. Gene set enrichment analysis: a knowledge-based approach for interpreting genome-wide expression profiles. Proc Natl Acad Sci U S A. 2005;102:15545–15550.

20. Liberzon A, Subramanian A, Pinchback R, Thorvaldsdottir H, Tamayo P, Mesirov JP. Molecular signatures database (MSigDB) 3.0. Bioinformatics. 2011;27:1739–1740.

21. Hafemeister C, Satija R. Normalization and variance stabilization of single-cell RNA-seq data using regularized negative binomial regression. Genome Biol. 2019;20:296.

22. Zappia L, Oshlack A. Clustering trees: a visualization for evaluating clusterings at multiple resolutions. Gigascience. 2018;7.

23. Yu G, Wang LG, Han Y, He QY. clusterProfiler: an R package for comparing biological themes among gene clusters. OMICS. 2012;16:284–287.

24. Roberts K, Aivazidis A, Kleshchevnikov V, Li T, Fropf R, Rhodes M, Beechem JM, Hemberg M, Bayraktar OA. Transcriptome-wide spatial RNA profiling maps the cellular architecture of the developing human neocortex. 2021.

25. Tarazona S, Garcia-Alcalde F, Dopazo J, Ferrer A, Conesa A. Differential expression in RNA-seq: a matter of depth. Genome Res. 2011;21:2213–2223.

26. NanoString Technologies I. MAN-10117-04 GeoMx - NGS Readout Library Prep User Manual. 2021.

27. Sequencing Requirements for Visium Spatial Gene Expression [https://www.10xgenomics.com/support/spatial-gene-expression-fresh-frozen/documentation/steps/sequencing/sequencing-requirements-for-visium-spatial-gene-expression]

28. Sequencing Requirements for Visium Spatial Gene Expression for FFPE [https://www.10xgenomics.com/support/spatial-gene-expression-ffpe/documentation/workflows/ffpe-v-1/steps/sequencing/sequencing-requirements-for-visium-spatial-gene-expression-for-ffpe]

29. Ni Z, Prasad A, Chen S, Halberg RB, Arkin LM, Drolet BA, Newton MA, Kendziorski C. SpotClean adjusts for spot swapping in spatial transcriptomics data. Nat Commun. 2022;13:2971.

30. Kim HK, Park KH, Kim Y, Park SE, Lee HS, Lim SW, Cho JH, Kim JY, Lee JE, Ahn JS, et al. Discordance of the PAM50 Intrinsic Subtypes Compared with Immunohistochemistry-Based Surrogate in Breast Cancer Patients: Potential Implication of Genomic Alterations of Discordance. Cancer Res Treat. 2019;51:737–747.

31. Parker JS, Mullins M, Cheang MC, Leung S, Voduc D, Vickery T, Davies S, Fauron C, He X, Hu Z, et al. Supervised risk predictor of breast cancer based on intrinsic subtypes. J Clin Oncol. 2009;27:1160–1167.

32. Hickey TE, Selth LA, Chia KM, Laven-Law G, Milioli HH, Roden D, Jindal S, Hui M, Finlay-Schultz J, Ebrahimie E, et al. The androgen receptor is a tumor suppressor in estrogen receptor-positive breast cancer. Nat Med. 2021;27:310–320.

33. Rodriques SG, Stickels RR, Goeva A, Martin CA, Murray E, Vanderburg CR, Welch J, Chen LM, Chen F, Macosko EZ. Slide-seq: A scalable technology for measuring genome-wide expression at high spatial resolution. Science. 2019;363:1463–1467.

34. Chen KH, Boettiger AN, Moffitt JR, Wang S, Zhuang X. RNA imaging. Spatially resolved, highly multiplexed RNA profiling in single cells. Science. 2015;348:aaa6090.

35. Lubeck E, Coskun AF, Zhiyentayev T, Ahmad M, Cai L. Single-cell in situ RNA profiling by sequential hybridization. Nat Methods. 2014;11:360–361.

36. Jerby-Arnon L, Neftel C, Shore ME, Weisman HR, Mathewson ND, McBride MJ, Haas B, Izar B, Volorio A, Boulay G, et al. Opposing immune and genetic mechanisms shape oncogenic programs in synovial sarcoma. Nat Med. 2021;27:289–300.

37. Meylan M, Petitprez F, Becht E, Bougouin A, Pupier G, Calvez A, Giglioli I, Verkarre V, Lacroix G, Verneau J, et al. Tertiary lymphoid structures generate and propagate anti-tumor antibody-producing plasma cells in renal cell cancer. Immunity. 2022;55:527–541 e525.

38. Erickson A, He M, Berglund E, Marklund M, Mirzazadeh R, Schultz N, Kvastad L, Andersson A, Bergenstrahle L, Bergenstrahle J, et al. Spatially resolved clonal copy number alterations in benign and malignant tissue. Nature. 2022;608:360–367.

39. Hudson WH, Sudmeier LJ. Localization of T cell clonotypes using the Visium spatial transcriptomics platform. STAR Protoc. 2022;3:101391.

40. Yang L, Yang Z, Danaher P, Zimmerman S, Hether T, Henderson D, Beechem J. Background modeling, Quality Control and Normalization for GeoMx RNA data with GeoDiff. 2022.

41. He S, Bhatt R, Brown C, Brown EA, Buhr DL, Chantranuvatana K, Danaher P, Dunaway D, Garrison RG, Geiss G, et al. High-Plex Multiomic Analysis in FFPE Tissue at Single-Cellular and Subcellular Resolution by Spatial Molecular Imaging. bioRxiv. 2022.

42. Janesick A, Shelansky R, Gottscho AD, Wagner F, Rouault M, Beliakoff G, de Oliveira MF, Kohlway A, Abousoud J, Morrison CA, et al. High resolution mapping of the breast cancer tumor microenvironment using integrated single cell, spatial and in situ analysis of FFPE tissue. bioRxiv. 2022.

