## Supplementary figures and images for "An experimental comparison of the Digital Spatial Profiling and Visium spatial transcriptomics technologies for cancer research"

### Supplemental Figure 1

#### Supplementary Figure 1

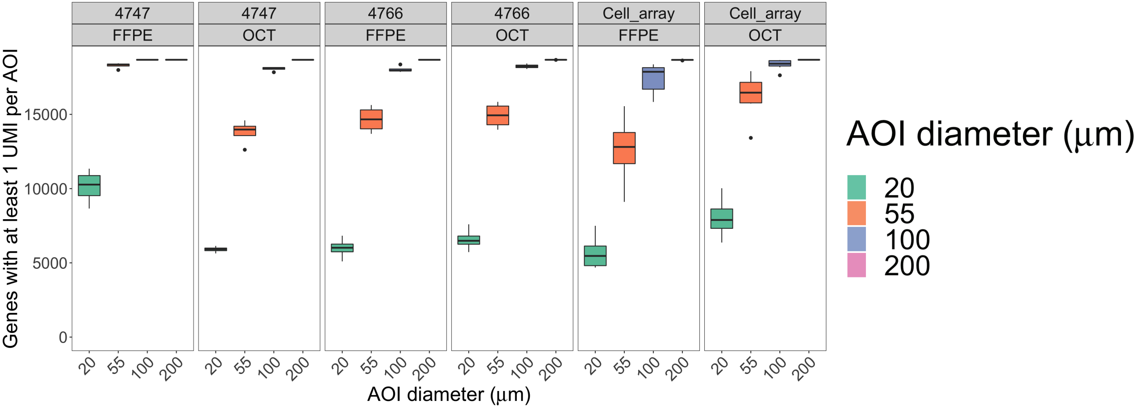

### Supplemental Figure 2

#### Supplementary Figure 2

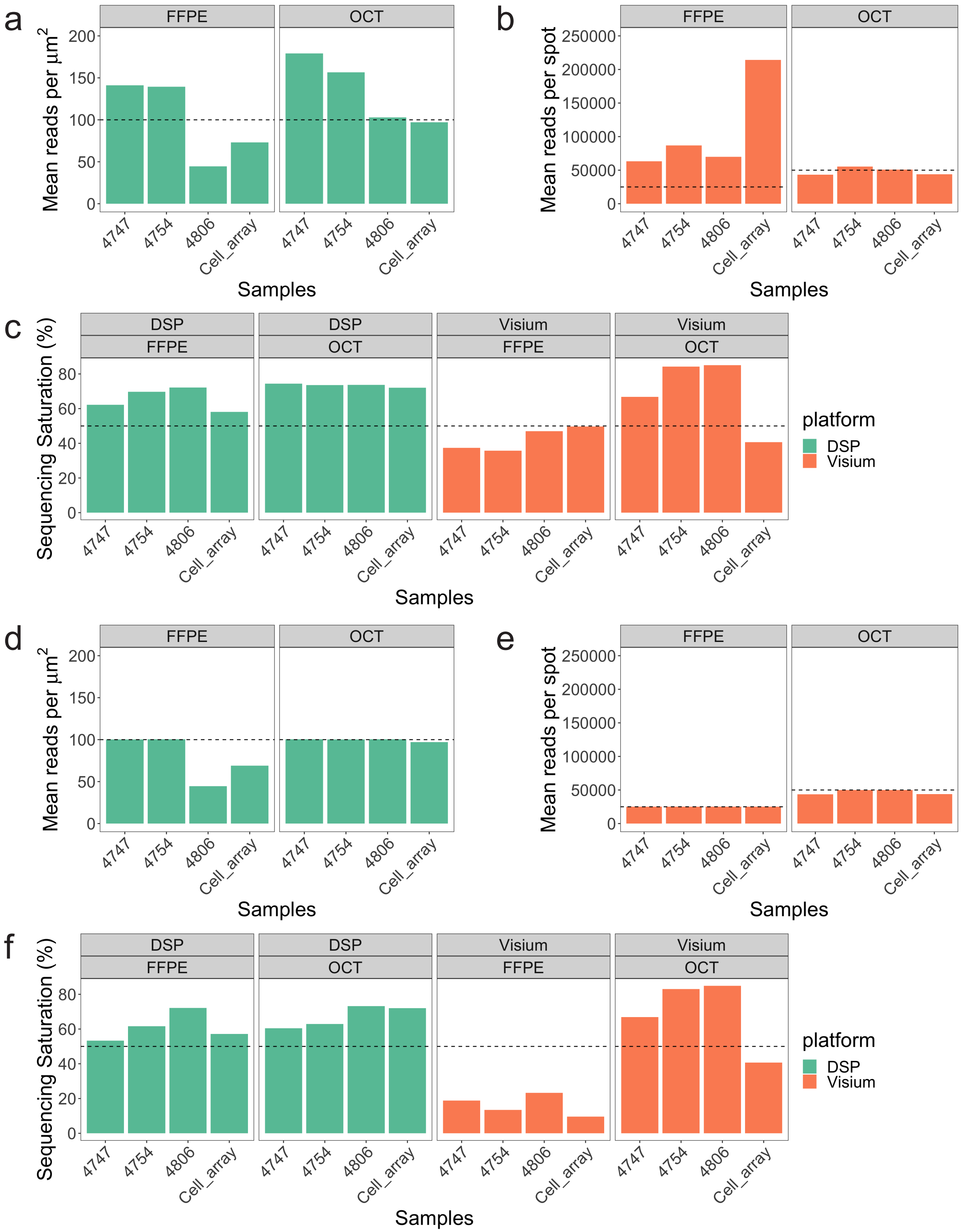

### Supplemental Figure 3

#### Supplementary Figure 3

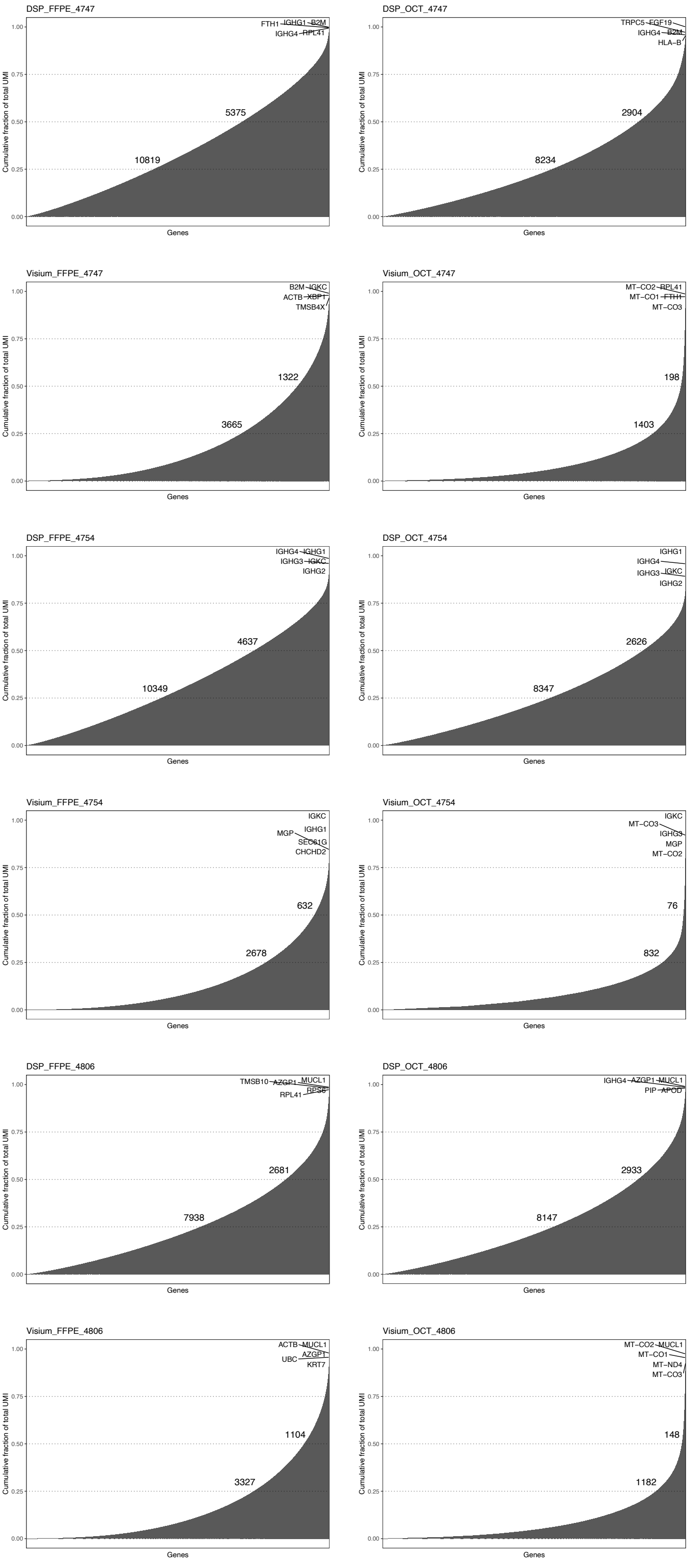

### Supplemental Figure 4

#### Supplementary Figure 4

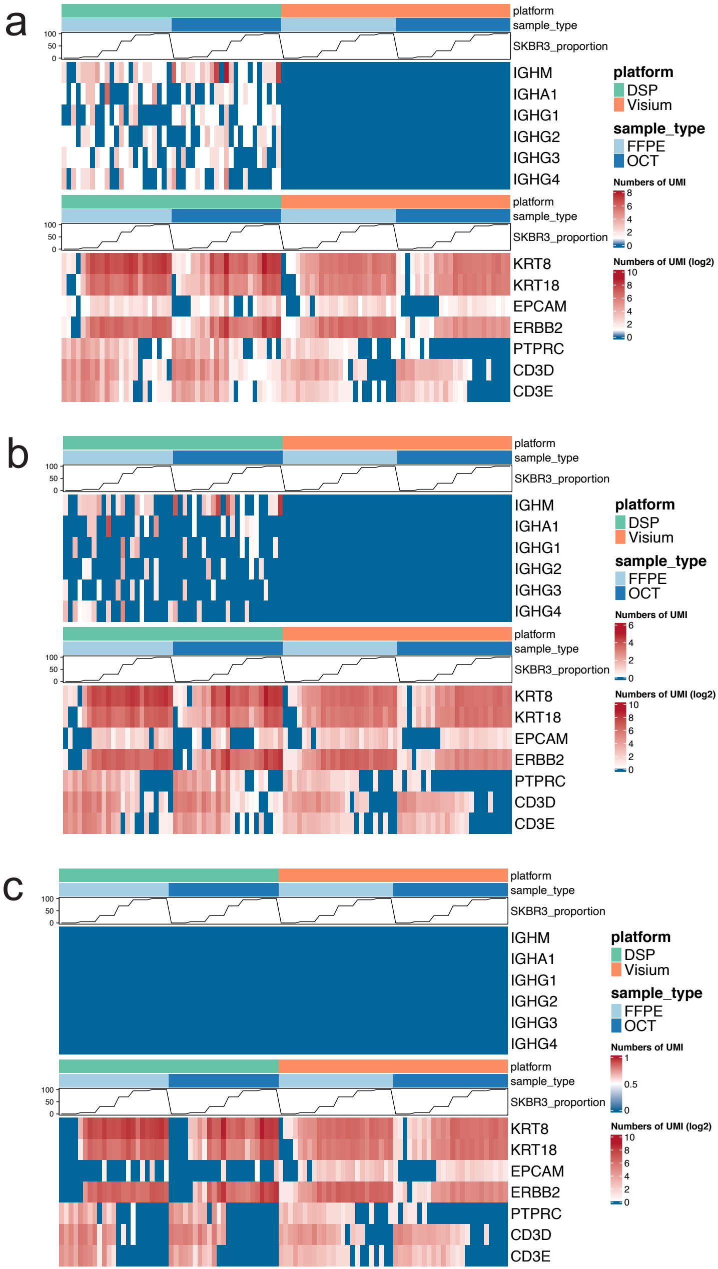

### Supplemental Figure 5

#### Supplementary Figure 5

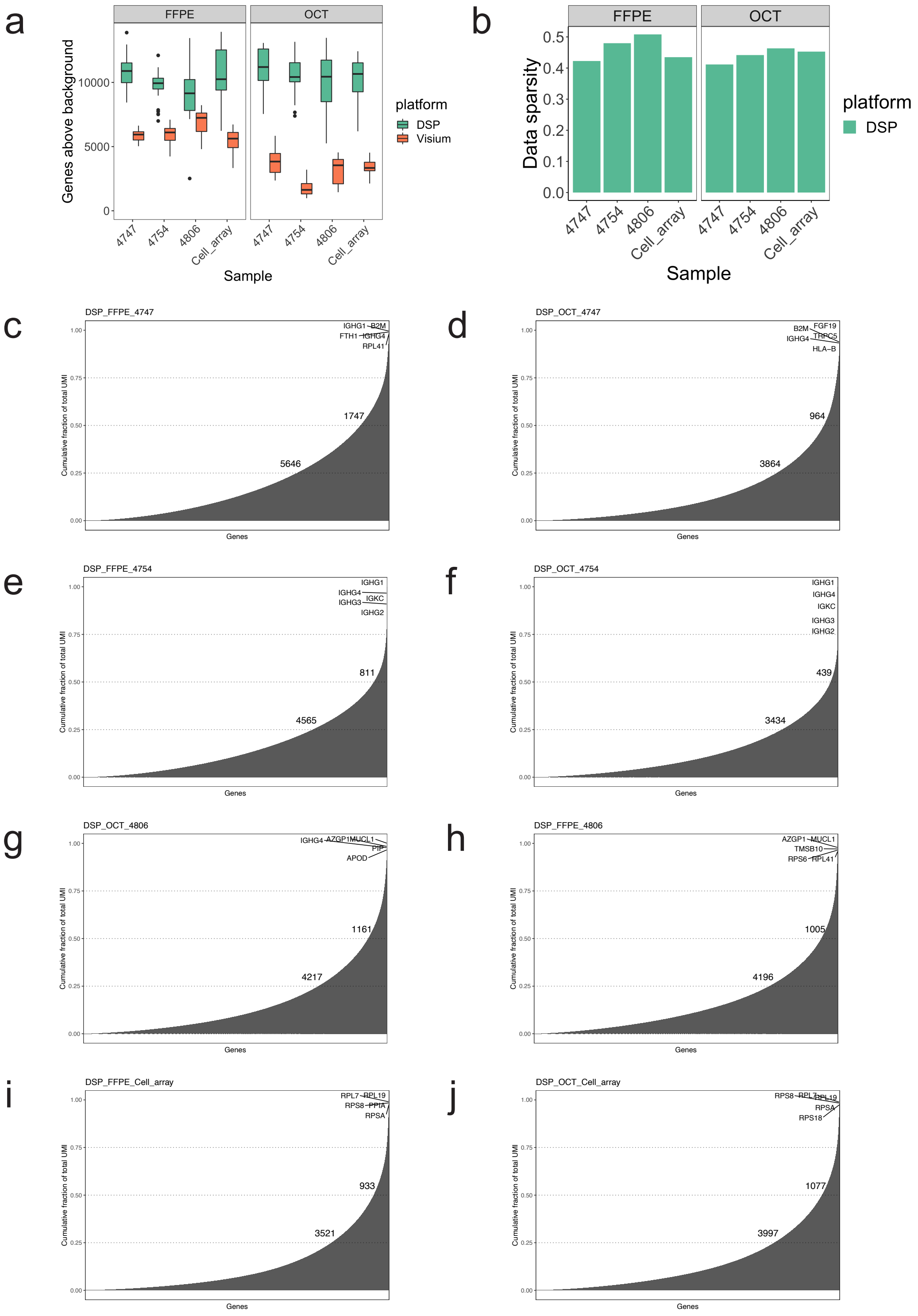

### Supplemental Figure 6

#### Supplementary Figure 6

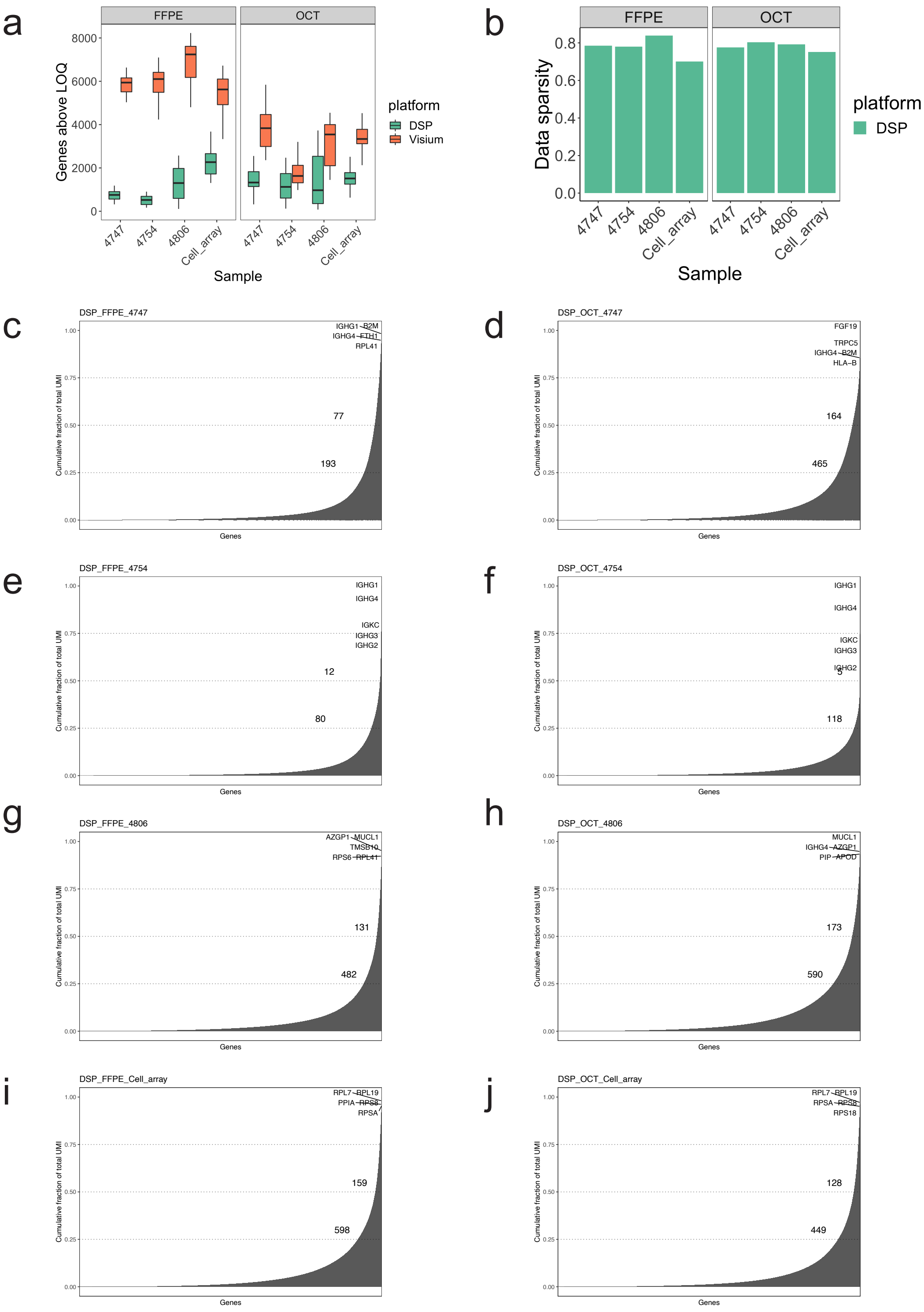

### Supplemental Figure 7

#### Supplementary Figure 7

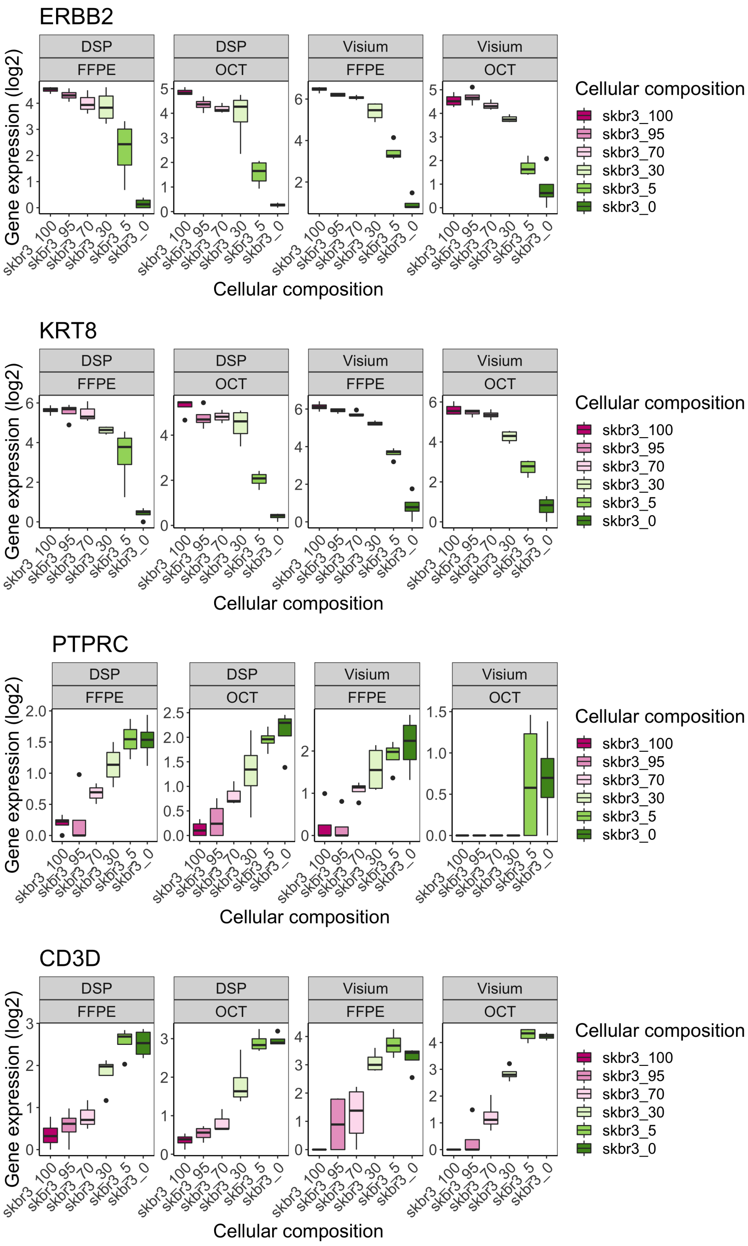

### Supplemental Figure 8

#### Supplementary Figure 8

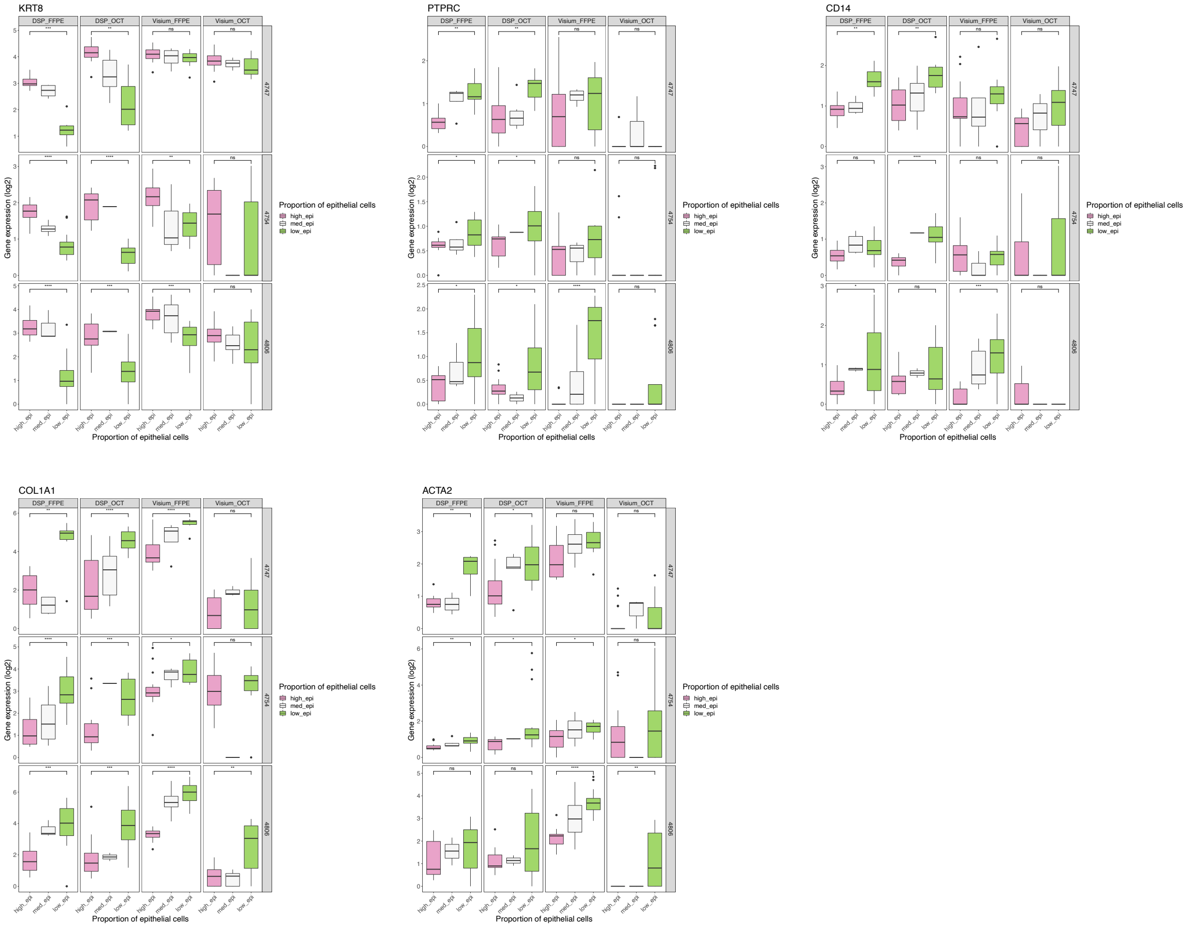

### Supplemental Figure 9

#### Supplementary Figure 9

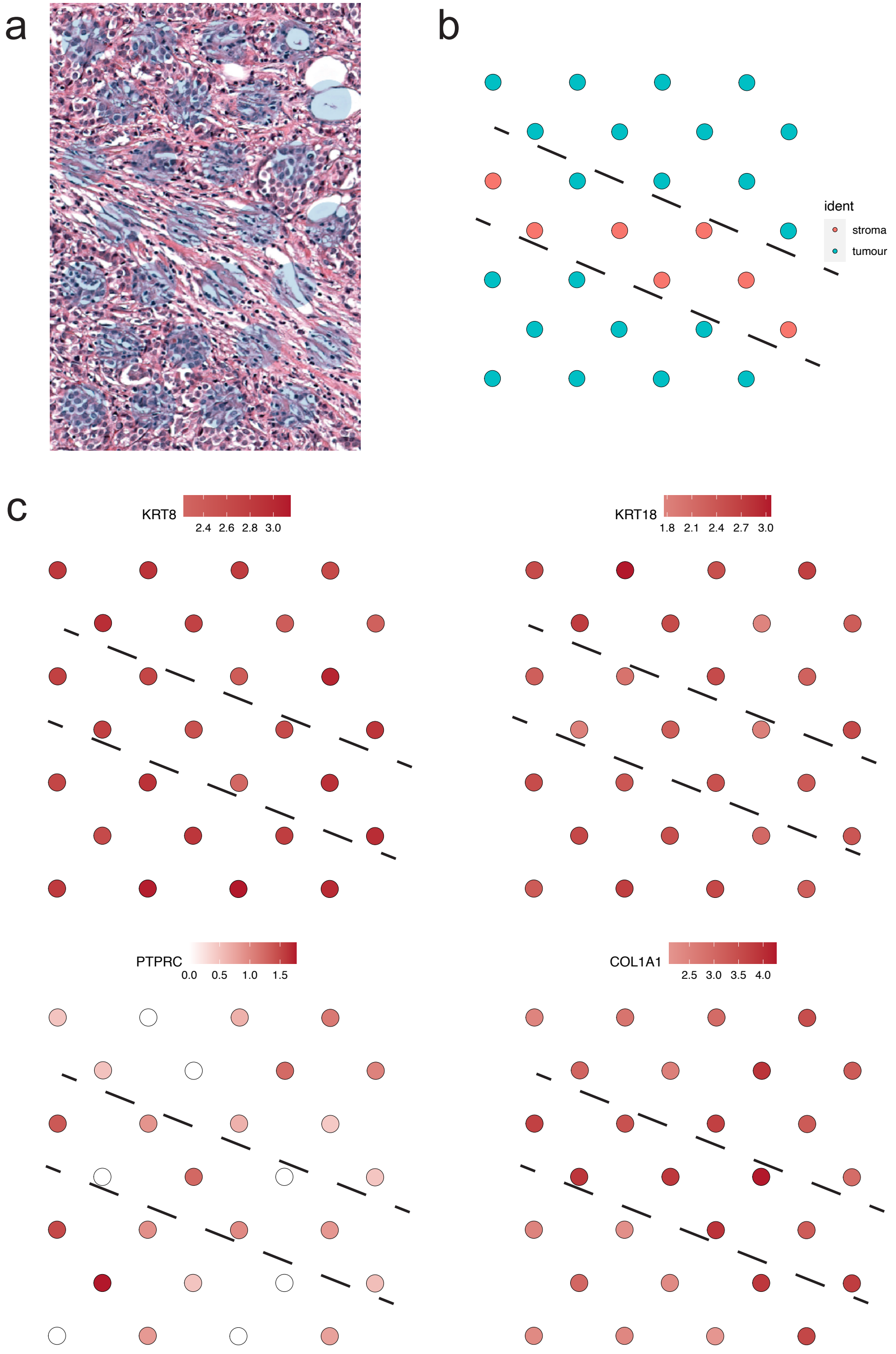

### Supplemental Figure 10

#### Supplementary Figure 10

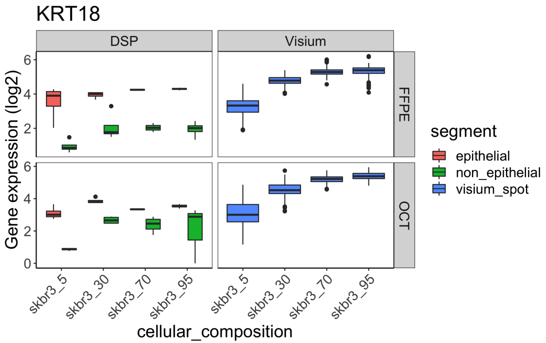


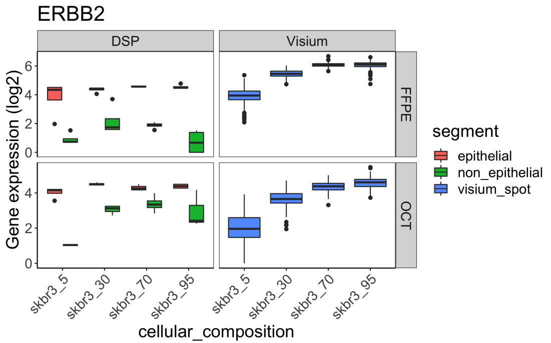


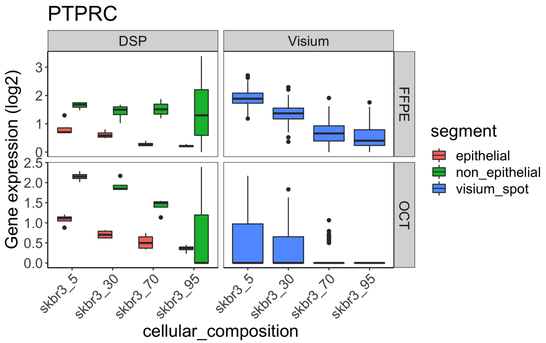


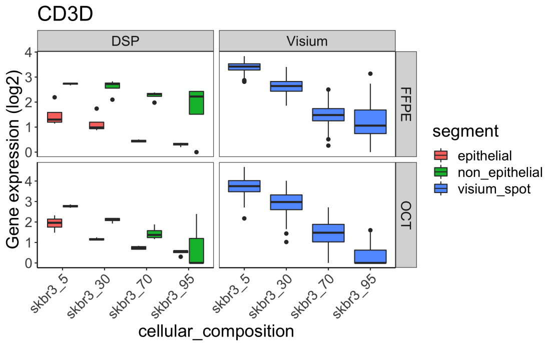

### Supplemental Figure 11

#### Supplementary Figure 11

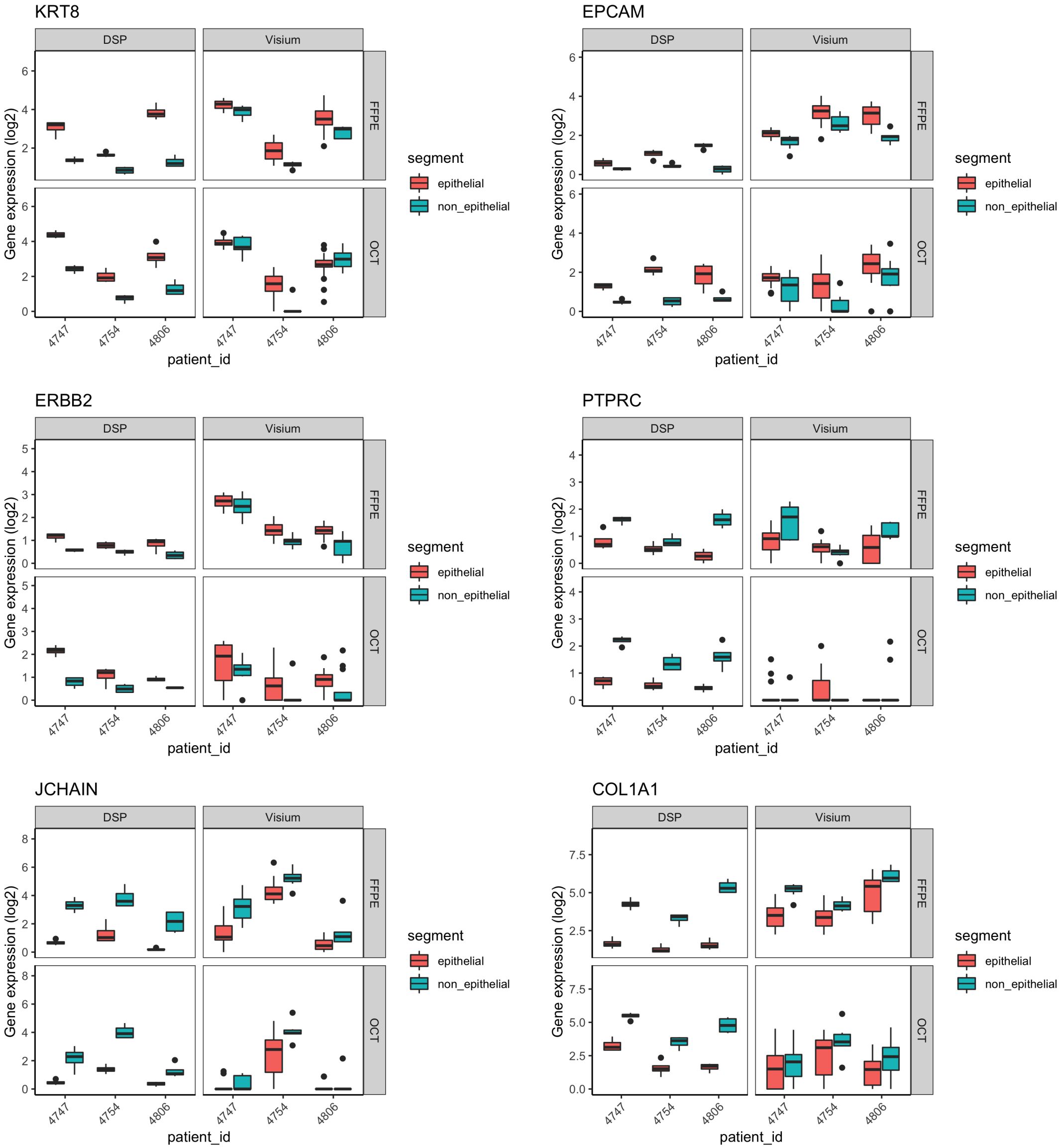

### Supplemental Figure 12

#### Supplementary Figure 12

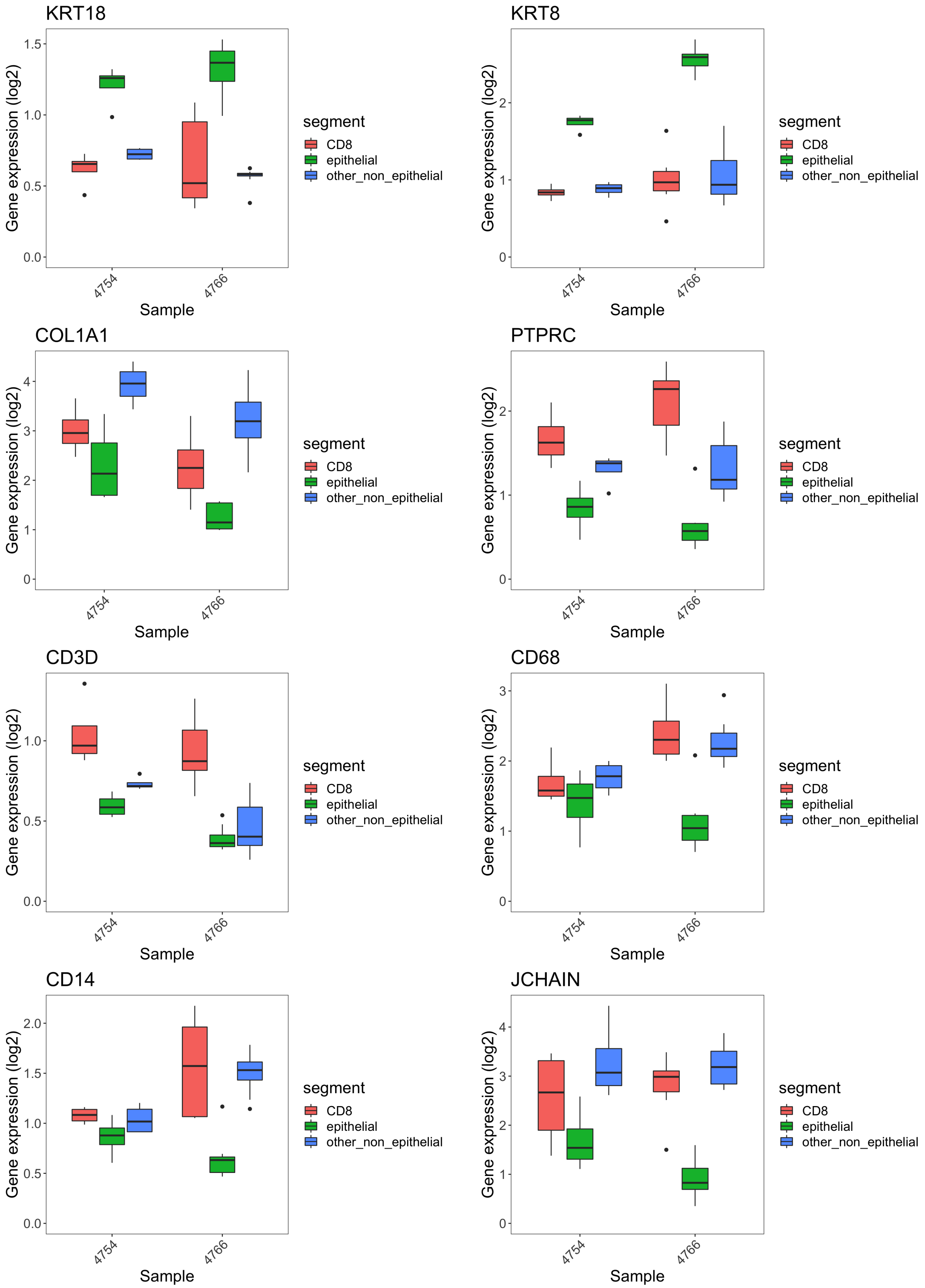

### Supplemental Figure 13

#### Supplementary Figure 13

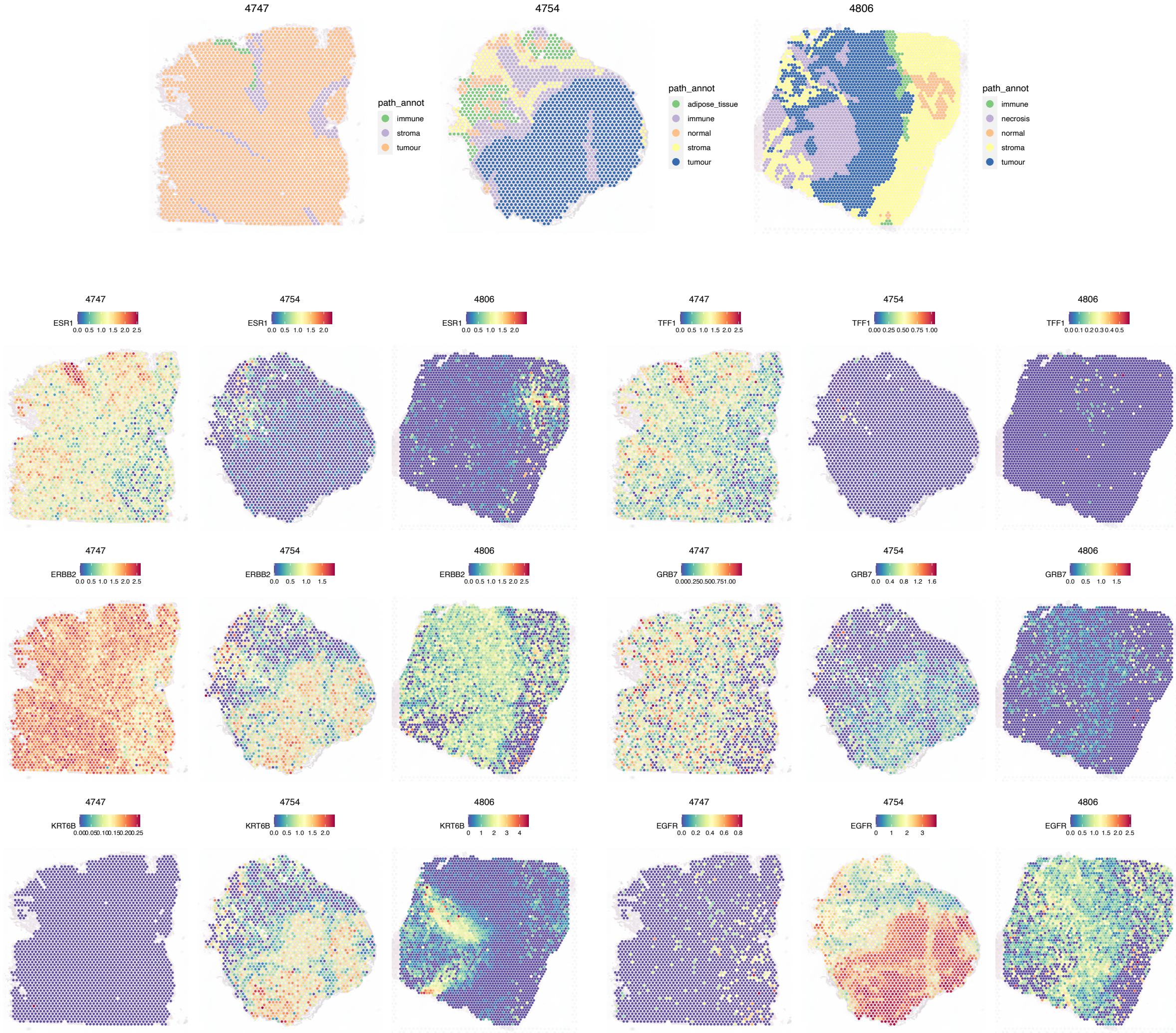

### Supplemental Figure 14

#### Supplementary Figure 14

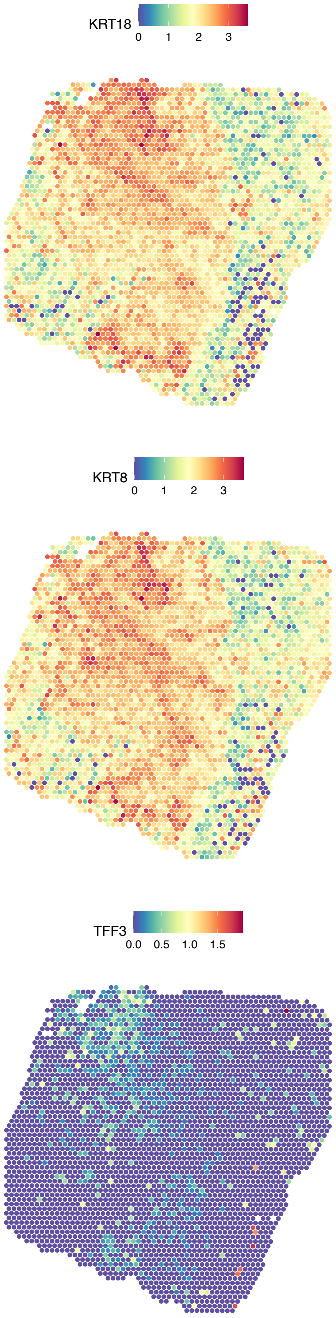

### Supplemental Figure 15

#### Supplementary Figure 15

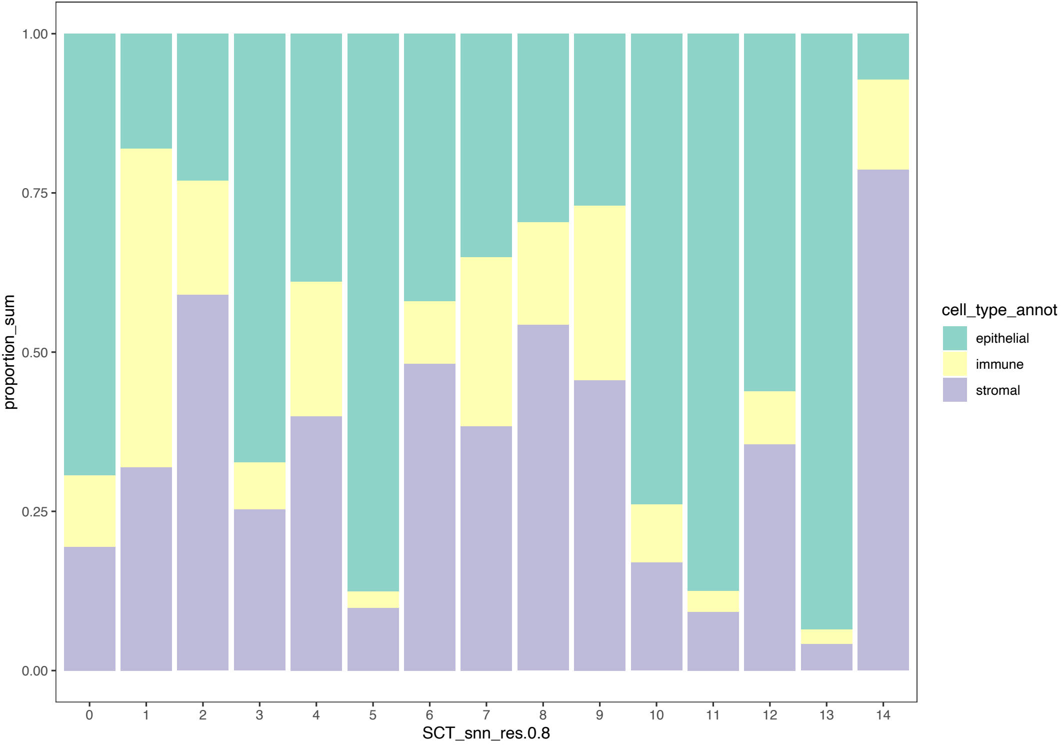

### Supplemental Figure 16

#### Supplementary Figure 16

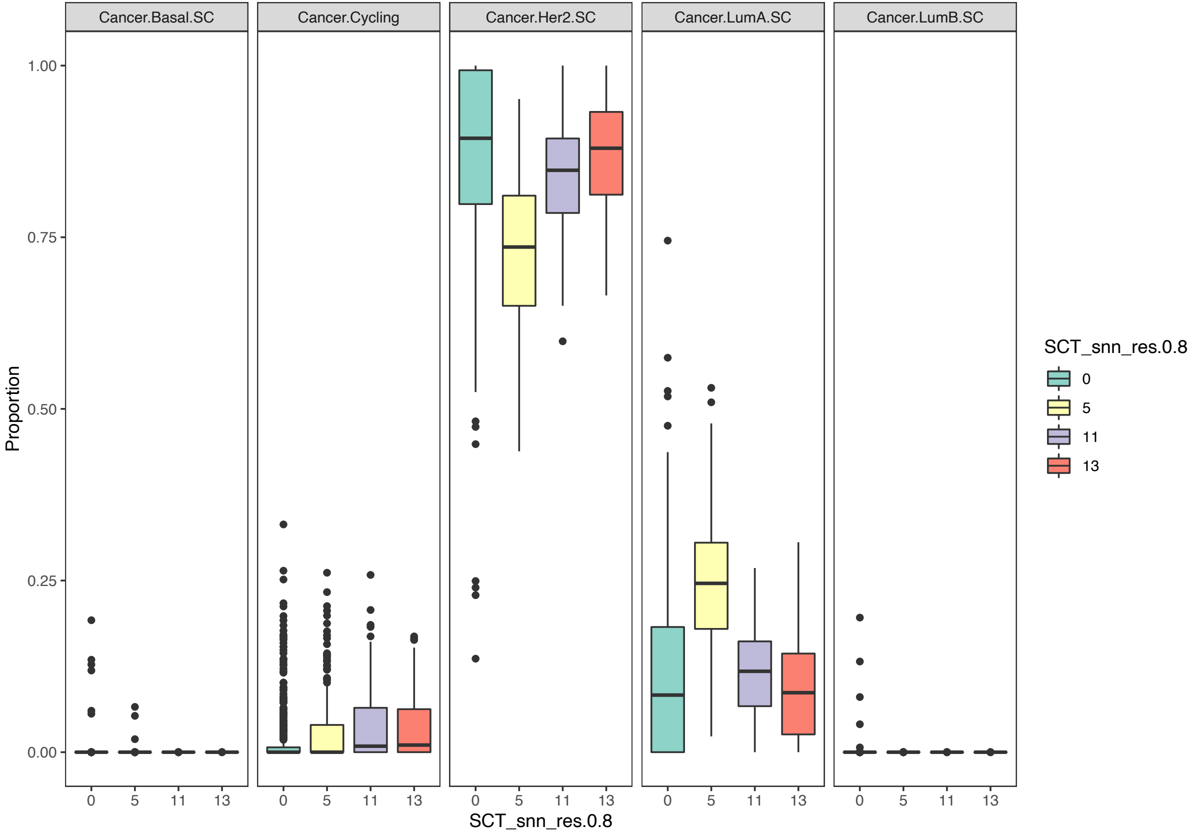
